# LARP1-DM15/Sgt is a reader for cap-adjacent 2’-*O*-ribose methylation in TOP ribosomal protein mRNAs required for localization to synapses

**DOI:** 10.64898/2026.09.17.752351

**Authors:** Yuan W. Tian, Yavor Hadzhiev, Giancarlo Abis, Martin Dodel, Emilie L. Alard, David W. J. Mcquarrie, Faraz K. Mardakheh, Maria R. Conte, Ferenc Müller, Matthias Soller

## Abstract

The most prominent mRNA modification in animals and many of their parasites is 2’-O-ribose methylation of cap-adjacent nucleotides (cOMe) introduced by cap-methyl transferases (CMTrs). How these modifications impact on gene expression and are decoded by reader proteins, however, remains uncertain. Through analysis of transcriptional start-site usage, we discovered a bias in ribosomal protein mRNAs starting with the Terminal Oligo Pyrimidine (TOP) motif, but not other TOP mRNAs in the absence of cOMe. cOMe stabilizes TOP mRNAs, enhances binding of the LARP1-DM15 domain to the TOP mRNA 5’ end and is assisted by the tetratricopeptide domain-containing protein Sgt/SGTA/B. This conserved complex is required for localization of TOP ribosomal protein mRNAs to *Drosophila* synapses. Our study reveals an mRNA methylation-mediated mechanism to maintain local protein synthesis remote from the nucleus for sustained synaptic functions.

## Introduction

Methylation of cap-adjacent nucleotides (cOMe) is an essential feature of animal messenger RNAs (mRNA) and those of many viruses and protists(Galloway and Cowling 2019; Lee et al. 2020; Haussmann et al. 2022; Anreiter et al. 2023; Despic and Jaffrey 2023; Dohnalkova et al. 2023; Clemens et al. 2025). Cap methyl transferases (CMTrs) add methylation at the 2’-*O*-ribose position of the first (cap1) and to a lesser extent also to the second cap-adjacent nucleotide (cap1/2)(Belanger et al. 2010; Werner et al. 2011; Dix et al. 2022; Haussmann et al. 2022; Anreiter et al. 2023). CMTr1 can methylate the first cap-adjacent nucleotide(Belanger et al. 2010; Dix et al. 2022). In contrast, CMTr2 is kinetically less active and methylates the second nucleotide, but in the absence of CMTr1 it can also methylate the first nucleotide in *Drosophila*(Dix et al. 2022). Despite its prevalence, the role of cOMe in regulating gene expression and whether it is decoded by methylation-sensitive reader proteins remains unresolved.

The first nucleotide of most mRNAs in *Drosophila*, and to a lesser extent in humans, is an adenosine followed by guanosine and uridine constituting a canonical consensus(Haussmann et al. 2022). In contrast, mRNAs of most cellular ribosomal proteins and many translation factors start with a cytosine followed by a number of uridines (Terminal Oligo Pyrimidine (TOP) motif)(Levy et al. 1991). The TOP motif is prominently recognized by the RNA binding protein LARP1 through its highly conserved C-terminal DM15 domain. Moreover, the DM15 domain binds to the cap *N7*-methylated guanosine and the cytosine in the first position to discriminate with high affinity against mRNAs with other first nucleotides(Fonseca et al. 2015; Lahr et al. 2017). CMTr1 is required for ribosomal protein gene expression(Liang et al. 2022), but whether cOMe impacts on LARP1-DM15 binding is not known. In addition to the DM15 domain, LARP1 contains the La-module consisting of an La motif and RNA recognition motif (RRM1) that binds other parts of the mRNA including the TOP motif and the polyA tail(Aoki et al. 2013; Al-Ashtal et al. 2021; Dock-Bregeon et al. 2021). LARP1 regulation of TOP mRNAs is directed by the mTOR pathway linking nutritional status to growth prominently dysregulated in many cancers(Thoreen et al. 2012; Saxton and Sabatini 2017).

While local translation of mRNAs into proteins is a prominent feature of neurons, it is generally employed by cells to organize translation of process specific genes in “translation factories”(Crawford et al. 2024). Local translation in dendrites employs multiple ribosomes (polysomes), while axonal and pre-synaptic translation occurs by a single ribosome (monosome) present on each mRNA(Biever et al. 2020). Ribosomes are assembled in the nucleolus and then transported along axons to synapses alongside many mRNAs for local translation. Unexpectedly, mRNAs coding for some ribosomal proteins (RPs) are also found at synapses(Shigeoka et al. 2019; Fusco et al. 2021; Yang et al. 2023). Notably, many of these synapse-localized RPs are predominantly positioned at the surface of the ribosome and local exchange in ribosomes assisted by dedicated chaperones may take place to repair ribosomes(Yang and Karbstein 2024). Since axonal nerve terminals can be far away from ribosome assembly sites in the nucleolus, such mechanism might be required to maintain local translation under high neuronal activity.

While in transit, ribosomes can translationally be made silent by specific RNA binding proteins including FMRP(Darnell et al. 2011; Kiebler and Bauer 2024). In addition, the nuclear cap binding complex has a stronger affinity for cOMe decorated mRNAs and is found at synapses alongside the Exon Junction Complex (EJC) deposited upon splicing in the nucleus(Haussmann et al. 2022). The EJC marks untranslated mRNAs as it is removed by the pioneer round of translation(Maquat et al. 2010; Topisirovic et al. 2011). Localization of FMRP and the EJC mark a pool of mRNAs for local translation and efficient localization of these mRNAs requires cOMe(Haussmann et al. 2022).

Both pre- and post-synaptic neuronal activity-dependent local translation are a hallmark of structural and functional plasticity linked to higher-order brain functions including learning and memory(Buffington et al. 2014; Holt et al. 2019; Bourke et al. 2023). In this context, cOMe devoid flies display many neurological phenotypes including reward learning deficits(Haussmann et al. 2022), but how cOMe contributes to local translation in synapses has not been determined.

Here we demonstrate that ribosomal protein-coding TOP mRNAs, but not other TOP mRNAs are downregulated in the absence of cOMe, that protects them from degradation. cOMe increases the affinity of the LARP1-DM15 domain to bind the cap and adjacent TOP motif, but not through recognition of cOMe. Here, through UV-crosslinking, we discovered a conserved reader for cOMe, the tetratricopeptide domain protein Sgt in *Drosophila* and SGTA and B in humans. Sgt together with cOMe can increase the affinity of DM15 by in the range of two orders of magnitude but does not bind RNA itself. Both LARP1-DM15 and Sgt mark TOP mRNAs for localization to synapses in the presence of cOMe.

## Results

### TOP ribosomal protein mRNAs are downregulated in the absence of cOMe in *Drosophila*

Since many genes have several transcription start sites (TSS) resulting in mRNA start nucleotide variation(Nepal et al. 2020; Anreiter et al. 2023; Wragg et al. 2023), we investigated whether cOMe impacts on TSS usage by performing CAGEseq in wild type and *CMTr1/2*^null^ mutant flies (*CMTr1* and *CMTr2* double knock-out)(Haussmann et al. 2022). We used the neuron-enriched head and thorax part because of increased expression of CMTrs in the nervous system(Haussmann et al. 2022). When analysing TSS upon CMTr1/2 loss, we noticed reduction of mRNAs starting with a C as first nucleotide (Fig 1A and S1A), confirming previous results(Liang et al. 2022). We performed an RNA-seq of wild type and *CMTr1/2*^null^ mutant flies and specifically analysed all TOP genes, and found predominantly ribosomal protein-coding genes to be significantly downregulated (Fig 1B). Nuclear encoded mitochondrial ribosomal proteins, whose mRNAs usually do not start with a TOP motif, were not significantly changed (Fig S1B). To validate downregulation of ribosomal protein genes, we performed qPCR for *RpL3,5,35* and *RpS16* genes. All four genes were significantly downregulated in head and thorax fraction of *CMTr1^null^*and *CMTr1/2^null^* mutants, but only partially in *RpS16* in CMTr2^null^ mutants indicating that cap1 is the predominant determinant of expression levels (Fig 1C).

**Figure 1.**
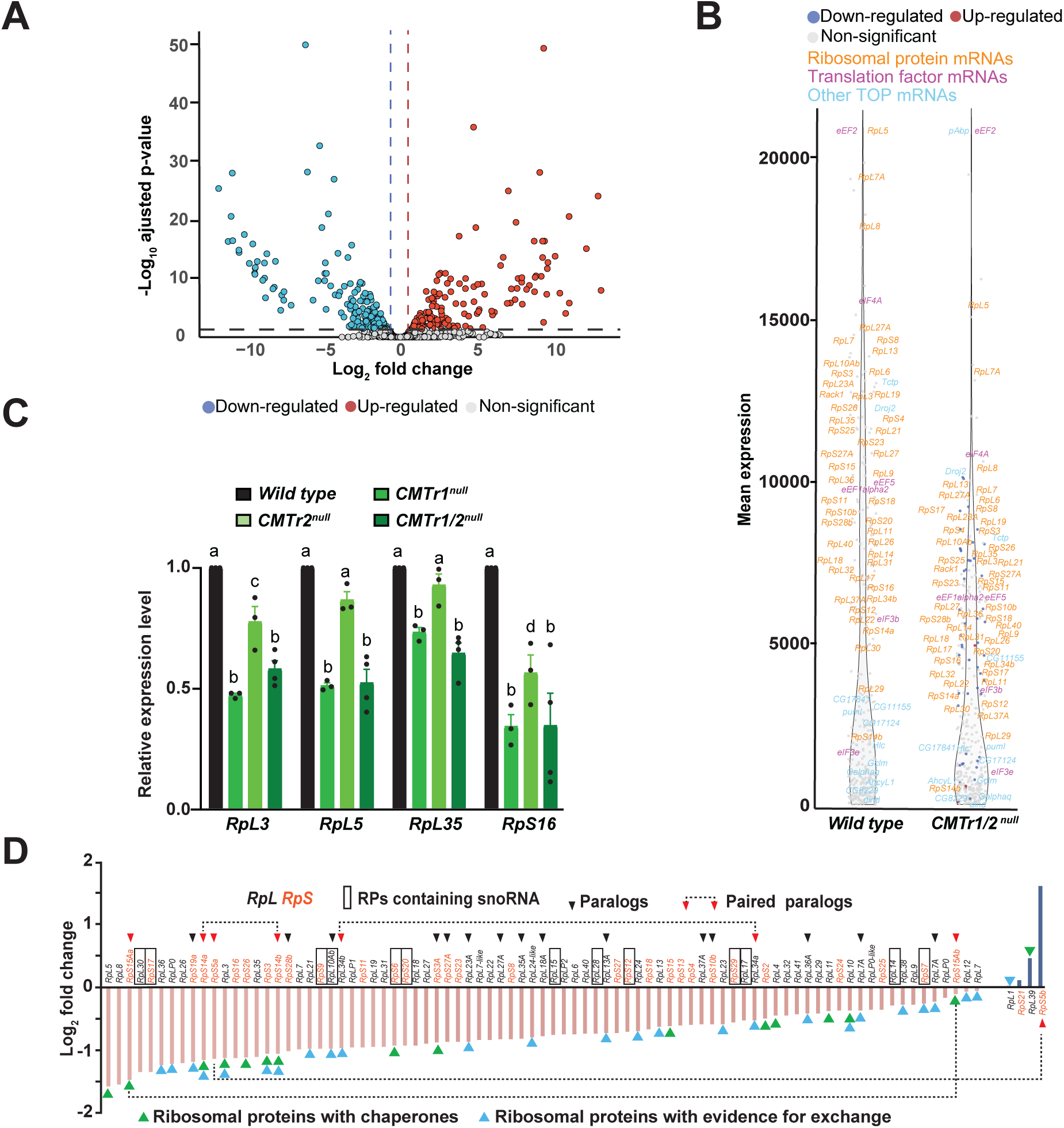
cOMe is required for ribosomal protein gene expression, but not other TOP mRNA genes. A) Volcano plot of CAGE-seq differential gene expression between wild type and *CMTr1/2^null^* (pink and blue dots: downregulated and upregulated mRNAs). B) Mean expression from RNA-seq of TOP mRNAs in wild type and *CMTr1/2^null^* mutant flies depicting expression levels of ribosomal protein mRNAs (orange), translation factor mRNAs (magenta), and other TOP mRNAs (blue). Significantly different expression is indicated by blue (down regulated) and red (upregulated) and non-significance by grey dots. C) Expression levels of *RpL3, RpL5, RpL35* and *RpS16* genes determined by RT-qPCR are shown as mean (n=3-4) with the standard error for wild type (black), *CMTr1^null^* (lime green), *CMTr2^null^* (olive green) and *CMTr1/2^null^* (avocado green). Statistically significant differences from ANOVA post-hoc comparisons are indicated by different letters (p≤0.003 except p≤0.017 for c). D) Graphic depiction of log_2_ fold changes in *CMTr1/2^null^* of ribosomal protein mRNAs from CAGE-seq (pink and blue bars: down-and up-regulated, black and orange: large and small ribosomal subunit.

Although not all TOP ribosomal protein mRNAs are equally downregulated, we did not observe a bias in affiliation to the ribosomal subunit or whether snoRNAs were present in the ribosomal protein genes (Fig 1D, Fig S2 and additional text). Notably, many of the encoded ribosomal proteins occupy the surface of ribosomes and can exchange with the assistance of certain chaperones (Fig 1E)(Yang and Karbstein 2024).

To determine differences at protein levels between wild type and *CMTr1/2*^null^ mutant flies, we use mass spectrometry. As a group, TOP ribosomal proteins are significantly downregulated in *CMTr1/2*^null^ mutants compared to wild type flies (p≤0.002, Fig S3). However, at the level of individual genes only RpS6 and RpLP2 were significantly downregulated pointing towards more specific roles in RNA localization and local translation.

### cOMe is required for TOP RNA stability

For mRNAs with G as a first nucleotide, cOMe protects against decapping by DXO in vitro and rescues *C. elegans CMTr1* mutant growth and fertility phenotypes (Picard-Jean et al. 2018; Clemens et al. 2025), but TOP ribosomal protein mRNA levels were not increased in *DXO^null^* mutant flies (Fig S4 and additional text). To evaluate whether cOMe stabilizes TOP motif RNA, we incubated m7G-capped RNA oligonucleotides with either cap0, cap1 or cap1/2 cOMe in nuclear and cytoplasmic HeLa or cytoplasmic *Drosophila* S2 cell extract (Fig 2A). After an initial drop of stability in the first 30 min, cap1 and to a stronger extent cap1/2 remained significantly more stable (Fig 2B-D).

**Figure 2.**
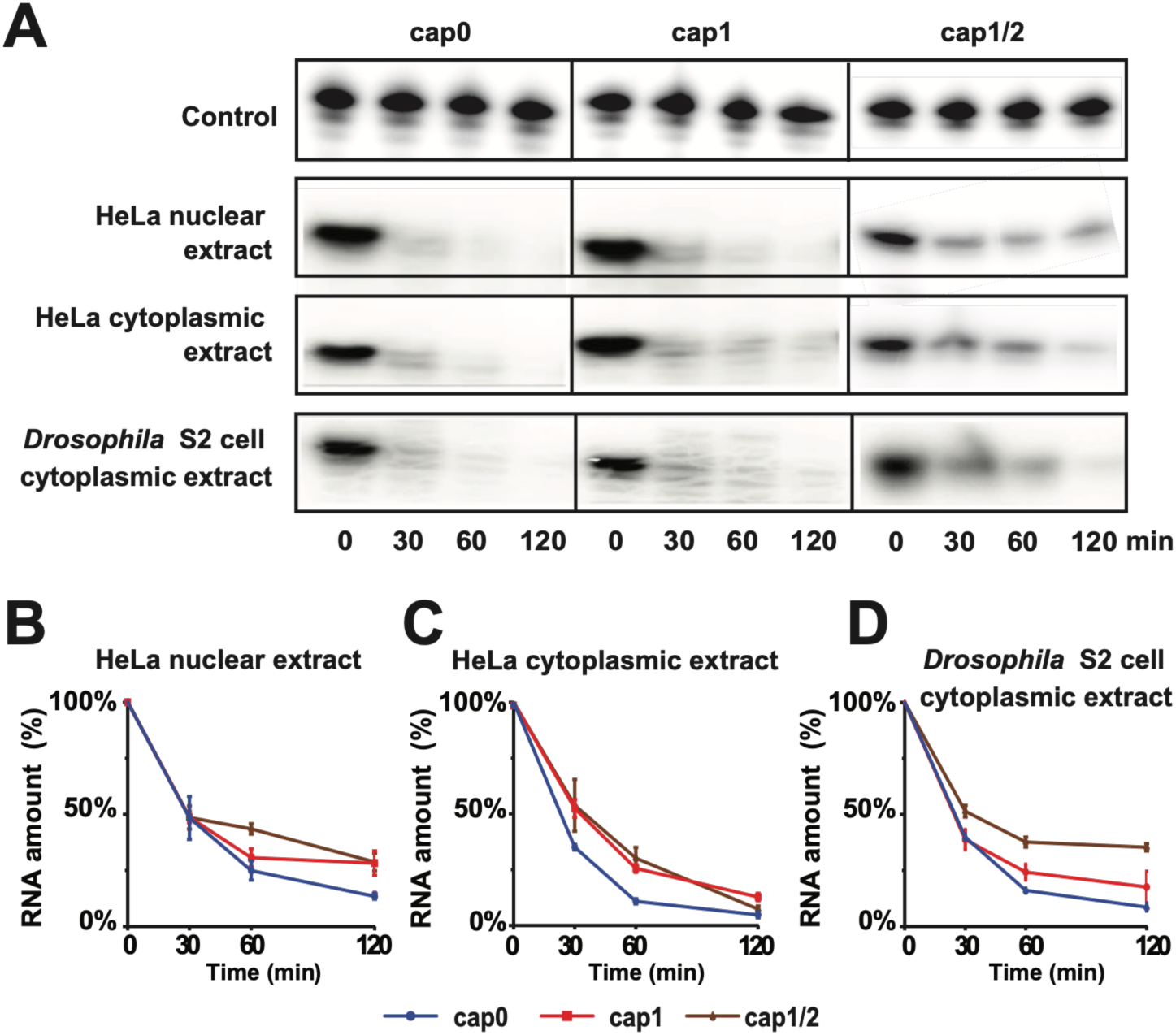
cOMe stabilizes TOP RNA in cellular extracts. A-D) Representative denaturing gels showing TOP RNA oligos (Rpl22, 25 nts) with varying methylation (cap0, cap1 and cap1/2) after incubation in cellular extracts (A) and quantification shown as mean (n=3) with the standard error for 1ap0 (blue), cap1 (red) and cap1/2 (brown) in HeLa nuclear (B) and cytoplasmic extract (C) and *Drosophila* S2 cell cytoplasmic extract (D).

### cOMe enhances binding of the LARP1-DM15 domain to TOP ribosomal protein mRNAs

A key feature of LARP1 is the C-terminal DM15 motif, which is highly conserved between *Drosophila* and humans and is absent in other LARPs (Fig 3A and Fig S5, and additional text)(Lahr et al. 2017; Nguyen et al. 2024). When we examined LARP1-bound mRNAs (LARP1-RIP-Seq data, GEO accession: GSE171350)(Martin et al. 2022) we observed a large overlap of genes differentially regulated in *CMTr1/2^null^* mutants (Fig 3B and C). To determine whether the DM15 domain regulates TOP mRNAs in vivo, we obtained a *LARP1* mutant lacking the DM15 domain (*LARP1^MI06928^* renamed to *LARP1^ΔDM15^*) and generated a *UAS LARP1-DM15* transgene for ectopic expression.

**Figure 3.**
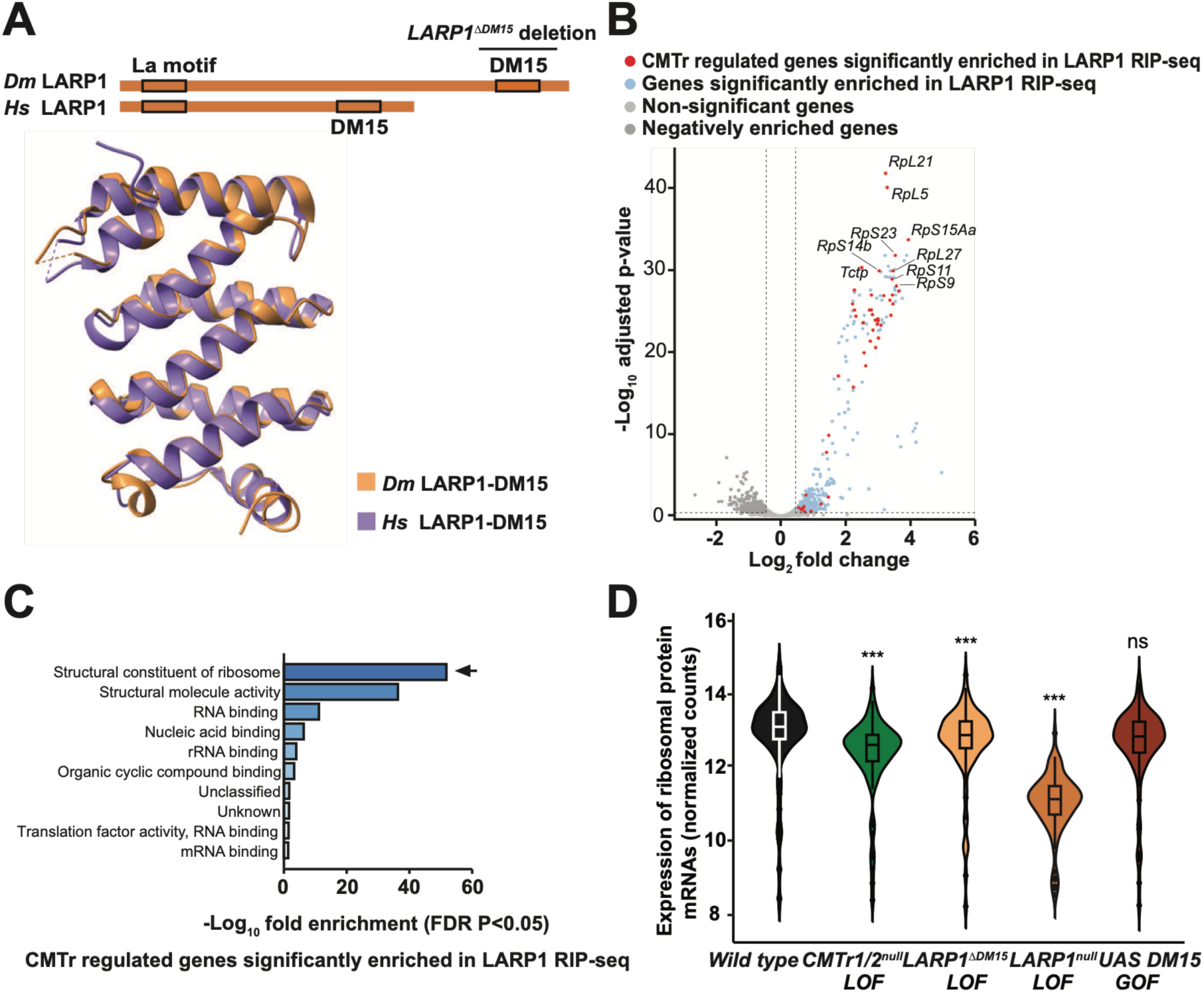
TOP ribosomal protein mRNAs are common targets of CMTrs and LARP1. A) Schematic of the *Drosophila* and human LARP1 domain structure and overlay of the *Drosophila* (brown) and human (purple) DM15 domain. B and C) Volcano plot of CAGE-Seq differential gene expression between wild type and *CMTr1/2^null^* superimposed on LARP1 targets determined by RIP-seq (B) and GO analysis (C). D) Violin plot depicting TOP ribosomal protein mRNA expression levels in neuron-enriched head/thorax of in wild type (black), *CMTr1/2^null^* (green), *LARP1^ΔDM15^* (almond), *LARP1^null^* (apricot) and *daGAL4 UAS DM15* (orange). Statistically significant differences to wild type from ANOVA post-hoc comparisons are indicated on top (***=p≤0.001).

Expression of ribosomal protein mRNAs was reduced in the DM15 deletion mutant (*LARP1^ΔDM15^*), but not as much as in *CMTr1/2^null^*or *LARP^null^* mutants (Fig 3D). Further expression of the LARP1-DM15 domain using neuronal *elavC155GAL4* did not increase ribosomal protein mRNA levels (Fig 3D). Intriguingly, in the *LARP1^ΔDM15^* mutant, we noticed a 2.2-fold upregulation suggesting compensatory regulation.

### Sgt binds to cOMe TOP RNA, forms a complex with LARP1 and is required for ribosomal protein mRNA expression

Although the LARP1-DM15 domain binds the cap guanosine moiety including the first C with a ∼100-fold higher affinity compared to capped RNA with a G at the first position(Lahr et al. 2017), the 2’position of ribose is facing outwards of the DM15 structure. To test whether binding of the DM15 domain is assisted by an additional reader, we employed UV-crosslinking in cellular extracts from HeLa and *Drosophila* cells using TOP and canonical start site (AGU) capped RNA oligo nucleotides with and without cOMe. These experiments identified a protein of the same size in human and *Drosophila* extracts (42 kDa) (Fig 4A). We purified this protein by immunoprecipitation of a terminally biotinylated RNA oligonucleotide and determined its identity by mass spectrometry as Small Glycine-rich Tetratricopeptide domain protein (Sgt in *Drosophila* and SGTA and B in humans, Fig 4B), which we validated by IP after UV-crosslinking in *Drosophila* extract with anti-Sgt antibody (Fig 4C). In this IP we also observed a larger protein of the size of LARP1. To validate that this Sgt interacting protein is LARP1, we did an IP after UV crosslinking in *Drosophila* extracts with an anti-LARP antibody and detected Sgt (Fig 4C). To validate that LARP1 interacts with Sgt in human cells, we did an IP after UV-crosslinking in HeLa extract with anti-LARP1 antibody and detected Sgt as well (Fig 4D).

**Figure 4.**
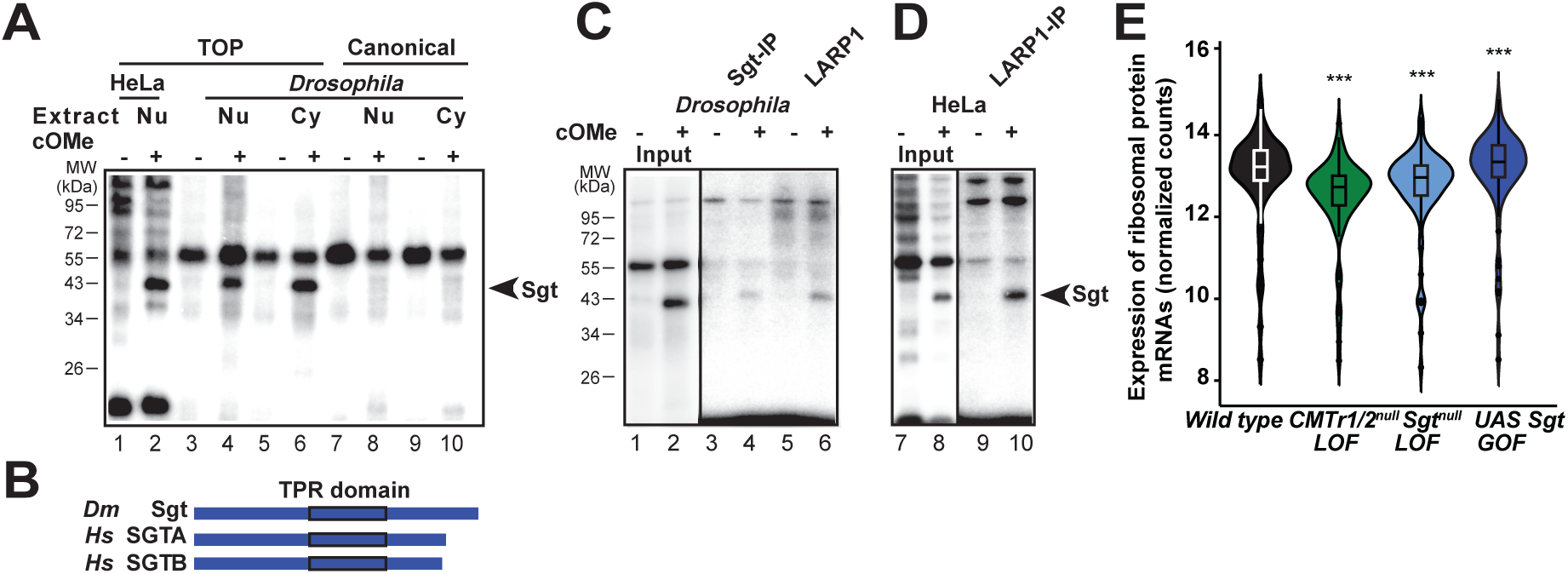
Sgt is a conserved reader for cOMe in TOP RNA and interacts with LARP1. A) UV-crosslinking of TOP (*Rpl22*) and canonical RNA (*per*) with or without cOMe after incubation in HeLa nuclear, and *Drosophila* S2 cytoplasmic and nuclear extracts. Molecular weight markers are shown on the left. B) Schematic of the *Drosophila* Sgt and human SGTA/B domain structure.| C) Immunoprecipitation with anti-Sgt (lanes 3 and 4) and anti-LARP1 antibodies (lanes 5 and 6) after UV-crosslinking in S2 cell extract of TOP RNA (*RpL22*) with or without cOMe in comparison to input (lanes 1 and 2). Molecular weight markers are shown on the left. D) Immunoprecipitation with anti-LARP1 antibodies (lanes 9 and 10) after UV-crosslinking in HeLa extract of TOP RNA (*RpL22*) with or without cOMe in comparison to input (lanes 7 and 8). E) Violin plot depicting TOP ribosomal protein mRNA expression levels in neuron-enriched head/thorax of wild type (black), *CMTr1/2^null^*(green), *Sgt^null^* (light blue) and *daGAL4 UAS Sgt* (blueberry). Statistically significant differences to wild type from ANOVA post-hoc comparisons are indicated on top (***=p≤0.001).

To test whether Sgt is indeed required for ribosomal protein mRNA expression in vivo, we obtained *Sgt^null^* mutants (*Sgt^56^*) (Uytterhoeven et al. 2015) and generated a UAS line for overexpression. *Sgt* like *CMTrs* and *LARP1* is broadly expressed (Fig S6A and B) and *Sgt^null^*mutants are viable as well.

In *Sgt^null^*mutants expression of ribosomal protein mRNAs is reduced and increases when *Sgt* is further expressed in neurons using *elavGAL4^c155^* compared to wild type flies (Fig 4E).

### Sgt enhances binding of LARP1-DM15 depending on cOMe

To examine the binding properties of the LARP1-DM15 domain and Sgt to capped and variable 2’-*O*-ribose methylated RNA oligos, we made recombinant protein in *E. coli* (Fig S7A) for in vitro binding assays and complex analysis using electrophoretic mobility shift assays (EMSAs)(Soller and White 2005; McQuarrie and Soller 2026). In these assays, LARP-DM15 bound cap0, cap1 and cap1/2 *RpL22* oligo with a Kd of 60±2.27 nM (n=3), 20±0.38 nM (n=3) 2±0.28 nM (n=3, Fig 5A-C and G), while the affinity of Sgt to bind these probes was above 10 µM (n=3).

**Figure 5.**
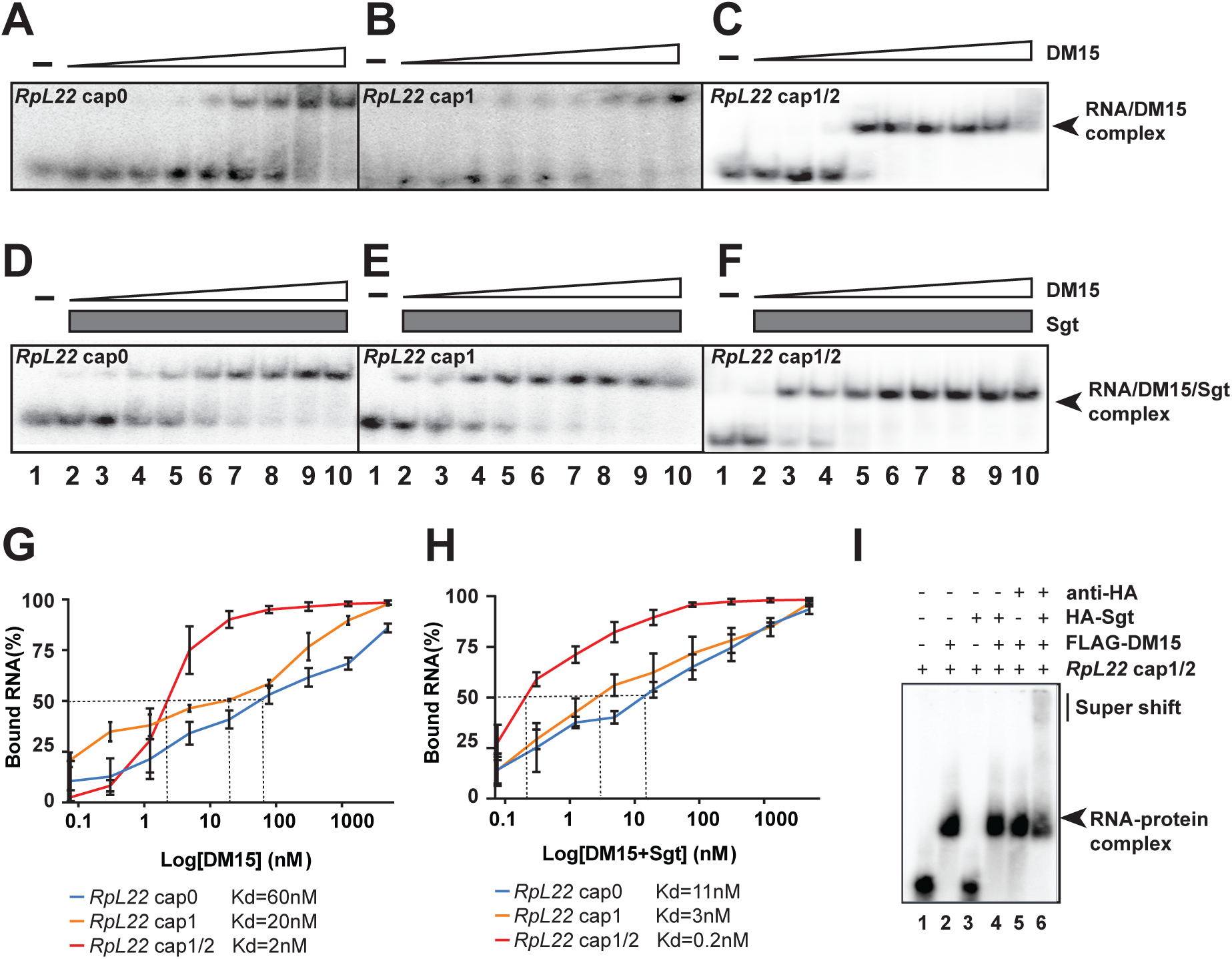
Binding of LARP1-DM15 to TOP RNA is enhanced by cOMe and Sgt. A-F) Electrophoretic mobility shift assays (EMSA) of various concentrations of DM15 (start from 5 µM, 4-fold serial dilution) and *RpL22* cap0, cap1 and cap1/2 RNA oligo (1 pM, A-C) alone or in the presence of Sgt (5 µM, D-E) and quantification thereof (G and H, cap0 in blue, cap1 in orange and cap1/2 in red). The mean and standard error of each concentration were calculated from three biological replicates, and Kd values are shown in G and H. I) EMSA of RNA alone (lane 1) and in the presence of DM15 (lane 2, with anti HA antibody lane 5), Sgt (lane 3) or both (lane 4) and super-shifted with anti HA antibody (lane 6).

To assess whether Sgt enhanced binding of LARP-DM15, we used a constant amount of Sgt in the same assay (5 µM). Addition of Sgt enhanced binding of LARP-DM15 to cap0, cap1 and cap1/2 *RpL22* oligo with a Kd of 11^±^0.79 nM (n=3), 3 ±0.29 nM (n=3) and 0.2±0.03 nM (n=3) with an overall 300-fold increase of affinity in the combined presence of cOMe (cap1/2) and Sgt (Fig 5D-F and H) but did not result in a more retarded complex in this native gel assay. To demonstrate presence of Sgt in the LARP-DM15 -RNA complex, we added an antibody to the binding mix resulting in a further shift of the complex (partial due to the limited amount of antibody available, Fig 5I).

To further validate enhanced binding of DM15 in the presence of cOMe and Sgt we repeated this experiment with *Rpl30* capped oligonucleotide which resulted in comparable affinities (Fig S7B-I). In addition, we performed binding assays using isothermal titration calorimetry (ITC), further confirming complex stoichiometry of 1:1:1 and an overall increase of LARP-DM15 affinity to capped TOP RNA of 36-fold in the presence of cOMe and Sgt (Fig S8).

### TOP mRNAs require cOMe for localization to synapses

We have previously shown that cOMe is required for localization of untranslated mRNAs to synapses and local translation(Haussmann et al. 2022). Since LARP1-DM15 and Sgt specifically bind to cOMe-containing TOP mRNAs, we determined whether they localize to synapses dependent on cOMe at third instar larval neuromuscular junctions (NMJs). Indeed, in the absence of cOMe, we find both DM15 and Sgt are essentially absent at synapses (Fig 6A). LARP1-DM15 is also absent in CMTr1 mutants which have little cap1/2 cOMe due to the presence of redundant CMTr2, or in the presence of only cap1 due to the absence of CMTr2. In contrast, Sgt is found at synapses in CMTr2 mutants which have cap1 mRNA, and at reduced levels in CMTr1 mutants, which have reduced levels of cap12 mRNA suggesting additional functions of Sgt(Uytterhoeven et al. 2015). In cell bodies, LARP1-DM15 localizes to the cytoplasm and cell membrane, while Sgt is strongly enriched at cell membranes (Fig S9A).

**Figure 6.**
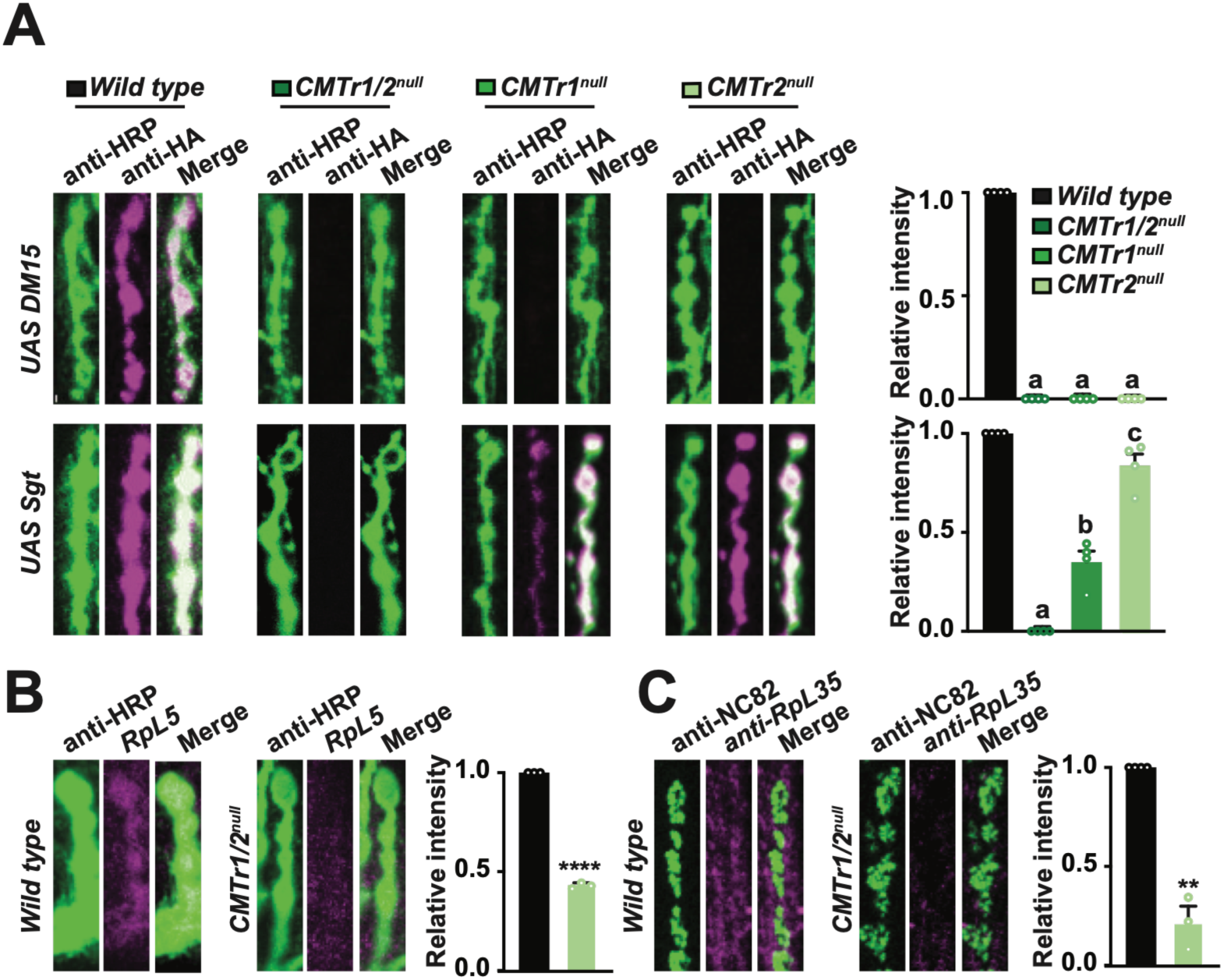
cOMe is required for TOP ribosomal protein mRNA and protein localization to synapses. A) Representative images from staining of synapses at third instar NMJs pre-synaptically expressing *UAS HA DM15* or *UAS HA Sgt* with *elavC155-GAL4* stained with an anti-HA antibody (magenta) and anti-HRP-antibodies (green) in wild type (black), *CMTr1^null^* (lime green), *CMTr2^null^* (olive green) and *CMTr1/2^null^* (avocado green) mutant larvae. The mean with the standard error of the intensity is shown on the right in arbitrary units. B and C) Representative images from RNA in situ hybridizations and anti-body stainings visualizing RpL35 RNA (B) and protein (C) in wild type and *CMTr1/2^null^*(avocado green) mutant larvae. NMJs were incubated with rabbit anti-HRP (green) and sheep anti-DIG-RpL35 antibody (B, magenta) or anti-RpL35 antibody (C, magenta). Mean intensity is normalized to wild type with standard error is shown on the right in arbitrary units.

Consistent with the absence of TOP mRNA localization at synapses in the absence of cOMe as marked with LARP1-DM15, levels of mRNA and protein for ribosomal protein RpL35 determined by RNA in situ hybridization and antibody staining, respectively, are also reduced (Fig 6B and C).

## Discussion

How the most prominent mRNA modification cOMe is decoded has remained enigmatic since its discovery almost 50 years ago(Furuichi et al. 1975; Wei et al. 1975). Here, we identified the first readers of this modification for TOP mRNAs, which start with C followed by several U’s. The cap and the first C are recognized by the DM15 domain of LARP1 and binding is enhanced by cOMe at the first nucleotide. Decoding of cOMe in TOP mRNAs further requires the TPR domain of the tetratricopeptide domain protein Sgt where cOMe enhances binding of the DM15/Sgt complex. The LARP1-DM15/Sgt forms a tight complex on the mRNA cap to protects TOP RNAs from degradation and to direct them to synapses.

### Distinct and overlapping roles for LARP1 and cOMe in regulating TOP mRNAs

A key role in TOP mRNA regulation is attributed to LARP proteins(Berman et al. 2021). Here, LARP1 is distinguished from LARP4, LARP6 and other LARP family members by the DM15 domain specifically recognizing the m7G cap and the first nucleotide, which must be a C. Loss of cOMe specifically reduced expression levels of TOP ribosomal protein mRNAs, but not other TOP mRNAs. This effect was partially recapitulated in the *LARP1^ΔDM15^* mutant, but we noted that the truncated *LARP1^ΔDM15^* transcript was upregulated. Likewise, TOP ribosomal protein mRNAs were further downregulated in *LARP1^null^*mutants compared to *CMTr1/2^null^* , and also reduced in *LARP4B^null^* and *LARP6^null^* mutants indicating additional parts of LARP proteins that can affect TOP ribosomal protein mRNA levels. In essence, this differential regulation suggests two pathways for regulating TOP ribosomal protein mRNA levels. In the first instance, cOMe increases the binding of LARP1-DM15 and Sgt to protect TOP ribosomal protein mRNAs from degradation. Secondly, binding of the La-module to the TOP motif, the body of the mRNA and the polyA tail results in stabilization, exemplified by LARP1 binding to mTOR mRNA (Mura et al. 2015; Al-Ashtal et al. 2021). Moreover, LARP6 has been shown to localize TOP ribosomal protein mRNAs to actin-rich cell protrusions for local translation(Dermit et al. 2020).

Expression of both CMTrs and LARP1 is enriched in the brain, but whether the reduction of TOP ribosomal proteins RNAs is neuronal, remains to be determined. Potentially, the fraction of TOP ribosomal proteins RNAs localizing to synapses could specifically be affected. A conserved feature of many ribosomal genes is the presence of dual start sites resulting in mRNAs with either a C or a purine as first nucleotide(Wragg et al. 2023). Although removal of CMTrs did not change start site use, mRNA fate could be directed by recruitment of mRNA degradation enzymes at specific promoters alongside co-transcriptionally recruited CMTr1.

### cOMe differentially stabilizes TOP mRNAs, but not canonical mRNAs

The start nucleotides of TOP mRNAs are distinct from the canonical AUG mRNA start, which is present in about 50% of mRNAs in *Drosophila*. Hence, LARP1-DM15 is specific to TOP mRNAs, but recognises only the first nucleotide. We identified Sgt through robust UV-crosslinking specifically to cOMe containing TOP RNA demonstrating close contact to RNA. However, the affinity for Sgt alone to bind RNA is below the threshold (>10 µM), but Sgt binds capped RNA in the presence of LARP1-DM15 and increases its affinity. Since DM15 covers the cytidine moiety of the first nucleotide entirely and is sensitive to cOMe, Sgt likely senses its methylation status and possibly recognizes the second nucleotide. However, further biophysical characterization is required to get insights DM15/Sgt/RNA complex architecture. Intriguingly, in both *Drosophila* and to a lesser extent in *C. elegans*, cOMe is primarily restricted to the first nucleotide(Dix et al. 2022). However, TOP mRNAs are only a minor fraction of all mRNAs. It is possible that specifically synaptic TOP mRNAs are cap1/2 marked, because cap1 is not sufficient to localize DM15 bound TOP mRNAs to synapses. In this sense, the methylation status might mark two pools of TOP mRNAs, one that is cap1 decorated for protection against degradation, and a second pool that is cap1/2 decorated to be localized to synapses. Since Sgt does localize to synapses when mRNA is only cap1 decorated, it could associate with another reader for the first nucleotide. In this case, however, stronger protein-protein interactions would need to compensate for loss of cOMe on the second nucleotide. Although Sgt has been shown to have other functions(Uytterhoeven et al. 2015), e.g. in membrane remodelling, this function also depends on cOMe, because Sgt is absent from synapses in the absence of cOMe.

Interferon induced transcripts (IFITs) are tetratricopeptide domain containing proteins, that can discriminate the methylation status of cap-adjacent nucleotides. In humans, there are five IFIT genes (IFIT1A, IFIT1B, IFIT2, IFIT3 and IFIT5), while mice have four (IFIT1, IFIT1b, IFIT1c, IFIT2 and IFIT3), but they have lost IFIT1A and duplicated IFIT1B twice(Mears and Sweeney 2018). Intriguingly, cOMe prevents binding of human IFIT1A, but will enhance binding of human IFIT1B and mouse IFIT1b(Kumar et al. 2014; Abbas et al. 2017; Mears and Sweeney 2018; Miedziak et al. 2020). Hence, cOMe can enhance binding of a reader on some transcripts, while in other situations cOMe prevents binding. However, the situation is more complicated as in humans IFIT3 acts as a co-factor of IFIT1A, while in mice IFIT1c is a co-factor of IFIT1 and IFIT1b(Mears and Sweeney 2018). How the two IFITs assemble to bind the mRNA cap and adjacent nucleotides as a complex remains to be determined, but could involve common principles also applicable to the DM15/Sgt complex.

LARP1-DM15 binding to TOP mRNAs is prominently regulated by mTOR phosphorylation to link nutrient status with cellular proliferation and growth(Thoreen et al. 2012; Saxton and Sabatini 2017; Goul et al. 2023). The low levels of cap2 in *Drosophila* and *C. elegans* could be linked to proliferation control as adult *Drosophila* and *C. elegans* used in previous studies have very little proliferating tissue. Since neurons are non-dividing cells cap1/2 can be adopted for localization of untranslated TOP mRNAs to synapses, while in proliferating cells cap1/2 could be dedicated to boost translation mediated by increasing ribosome numbers to promote proliferation and growth(Gao et al. 2026).

### Diversified local translation at pre-synaptic nerve terminals

Regulation of local translation at synapses comes in distinct flavours(Perez et al. 2021; Bourke et al. 2023; Cagnetta et al. 2023; Holt 2024). A main distinction is whether local translation occurs pre- or post-synaptic. Post-synaptic translation takes place in polysomes, while pre-synaptic translation occurs through monosomes, a single ribosome translating one mRNA. Ribosome are assembled in the nucleolus and then transported to synapses, which in large animals can be meters away from cell bodies. However, neurons can also do the final steps of ribosome assembly in axons(Fusco et al. 2025). The LARP1-DM15 domain is required to localize TOP ribosomal proteins to synapses. It is thus conceivable, that locally provided ribosomal proteins derived from synapse localized mRNAs could support ribosome function in local translation by local assembly to make new ribosomes, by providing specialized ribosomes or for repair. Exchange of damaged proteins in the ribosome has been observed after oxidative damage or growth factor stimulation(Shigeoka et al. 2019; Fusco et al. 2021; Yang and Karbstein 2024).

Loss of individual ribosomal proteins results in distinct phenotypes described as ribosomopathies(Mills and Green 2017), but differential expression of ribosomal proteins can also lead to ribosomes with specialised functions(Genuth and Barna 2018) resulting in adaptations to different cellular conditions(Ferretti et al. 2017; Cheng et al. 2019; Chen et al. 2026). CMTr1/2 loss result in many down-regulated TOP ribosomal protein RNAs arguing for a more global role in shaping neuronal function as indicated by distinct ribosomal protein mRNA expression profiles between excitatory and inhibitory neurons(Garat et al. 2026).

Alternatively, LARP1 binds TOP mRNAs together with ribosomal subunits and could substantially contribute to synapse localization(Gentilella et al. 2017; Fuentes et al. 2021; Saba et al. 2024). Because ribosomal protein mRNAs are among the most abundant transcripts in the cell, it might provide efficient means to transport ribosomes to synapses, which are arrested as monosomes on single mRNAs(Biever et al. 2020; Castillo et al. 2023).

Signalling by mTOR (target of rapamycin) kinase occurs through the two main multiprotein complexes mTORC1 and mTORC2 to regulate cell growth and proliferation. Intricate regulation adapts expression of TOP mRNAs to different cellular conditions including maintenance of basal translation under starvation(Gentilella et al. 2017; Fuentes et al. 2021; Schneider et al. 2022), but ribosomes can also be redirected by mTOR from TOP mRNAs for translation of essential mRNAs under severe stress(Fan et al. 2026). In neurons, however, mTOR regulates synaptic transmission by tuning the strength of synaptic connections. In particular, in mouse glutamatergic hippocampal neurons, the effects of mTORC1 are postsynaptic, while mTORC2 acts pre-synaptically(McCabe et al. 2020). In *Drosophila*, TOR is required for the retrograde regulation of synaptic homeostasis at third instar NMJs(Penney et al. 2012). Here, we discovered a role for mTOR target LARP1 in the regulation of local translation of ribosomal proteins, but how local translation will impact on neuronal plasticity through the regulation of local translation requires to be determined.

Another important role of mTOR signalling is in the regulation of autophagy which is also mediated by mTORC1 and 2 complexes and involves extensive membrane remodelling (Sun et al. 2023). Engulfment of proteins by the late endosomal membrane in a process called micro-autophagy is mediated by the chaperone Hsc70-4 facilitating delivery of targeted proteins to lysosomes(Uytterhoeven et al. 2015). Intriguingly, one of the most abundant synaptic proteins, Hsc70-4, is assisted by Sgt in *Drosophila* to support synaptic functions. Whether this role is associated with directing proteins to lysosomes is not clear in light of a role of lysosomes in transporting mRNA(De Pace et al. 2024). Although Sgt’s synaptic localization is dependent on cap12 decorated TOP mRNA localization, cap1 is sufficient to localize Sgt, but not DM15 to synapses. It is possible, that cap1 decorated Sgt mRNA can localize to synapses, but to rigorously test this hypothesis a neuron specific reporter driven by the endogenous promoter is required. In light of such scenario, Sgt might also exert functions independent of binding to RNA at synapses.

Taken together, our data suggest a mechanistic model for cOMe regulation of TOP mRNAs (Fig 7). In cell bodies, cap1’s main role is to protect TOP mRNAs from degradation. For TOP mRNAs to localize to synapses, however, cap1/2 is required as cap1 is not sufficient to localize Sgt to synapses. TOP mRNAs decorated with cap1/2 will then localize to synapses bound by LARP1-DM15 and Sgt, but to determine whether phosphorylation of LARP1 by TOR initiates translation requires endogenous promoter-based reporter transgenes.

**Figure 7.**
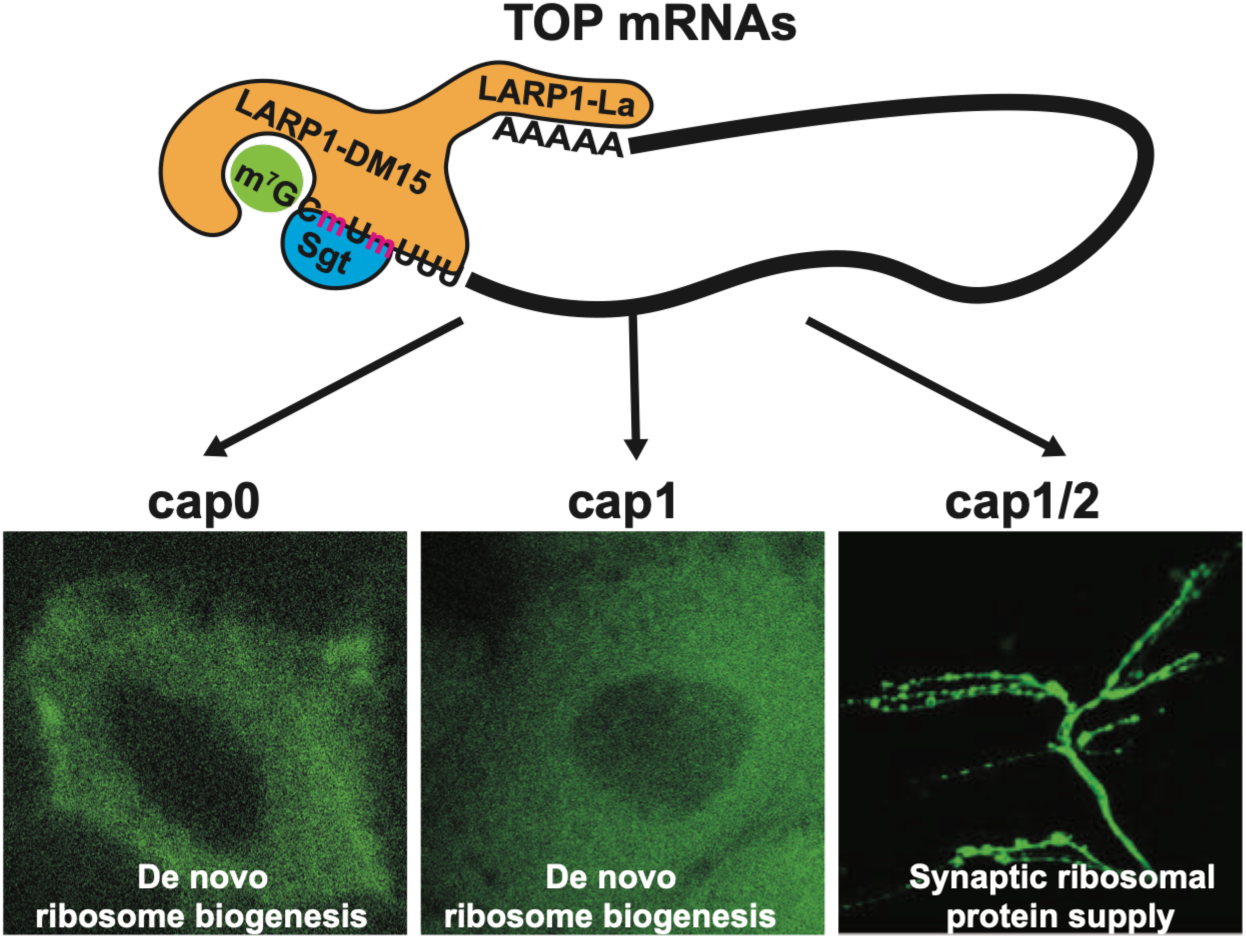
LARP1-DM15 is required for localization of TOP ribosomal protein mRNAs to synapses. Mechanistic model of cOMe decoding in TOP ribosomal protein mRNA by an DM15/Sgt reader complex.

Much of how local translation at synapses is regulated remains little understood due to many technical challenges. Given the prevalence of intellectual disability and neuronal disorder phenotypes associated with RNA modification pathways and LARP1 haploinsufficiency(Dezi et al. 2016; Chettle et al. 2024; Delaunay et al. 2024), key roles in the regulation of synaptic plasticity can be attributed to local translation that are sensitive to dosage. Expression of ribosomal proteins remote from the cell body points towards existence of self-sustained translation machinery essential for neuronal function in the context of neuronal activity by synaptic expression of proteins.

## Materials and METHODS

### Fly husbandry

All *Drosophila melanogaster* strains were reared at 25°C and 40%–60% humidity on standard cornmeal-agar food in a 12:12 h light: dark cycle. 14-18 h old embryos, wandering third instar larvae and 2-10 day old adult flies were used. Canton-S flies marked with y w were used as the wild type.

### Generation of recombinant DNA, transgenic fly strains and fly whole genome sequencing

*Drosophila LARP1-DM15* (amino acids 796-947) and *Sgt* (amino acids 1-331) were PCR amplified from cDNA made with Superscript II (Invitrogen) and cloned into NotI and XbaI cut modified *pUAST* vector containing an N-terminal HA and an *attB* site or Nhel and Spel cut *pUC 3GLA UAS* vector(Zaharieva et al. 2015; Haussmann et al. 2019; Haussmann 2024). Transgenic flies were then generated by phiC31 mediated integration into attP40.

The *Sgt^56^* allele(Uytterhoeven et al. 2015) was validated by whole genome sequencing as described(Lassota et al. 2025).

### CAGE Sequencing

To enrich for neurons, CAGE-Seq was done as previously described using Trizol extracted total RNA from 15 female *Drosophila* wild-type and *CMTr1/2*^null^ head and thoraxes(Haussmann et al. 2011; Wragg et al. 2023). CAGE-Seq data was analysed by the Bioconductor package CAGEr(Haberle et al. 2015).

### Proteomics

Protein lysates were prepared from 200 female *Drosophila* wild-type and *CMTr1/2*^null^ heads and thoraxes in 4% SDS, 50 mM Tris HCl pH 7.5, solubilised by extensive sonication, before being subjected to Trypsin digestion using Filter Aided Sample Preparation (FASP)(Wiśniewski et al. 2009), with modifications. Briefly, equivalent of 10 µg of lysate proteins were reduced with 10 mM DTT for 30 min at 25°C, and alkylated with 55 mM Iodoacetamide for 30 min at 25°C while protected from light. Samples were then diluted 8 fold by addition of UA buffer (8 M Urea, 100 mM Tris-HCl pH 8.5) and concentrated on a Vivacon 500, 30000 MWCO Hydrosart filter (Sartorius) by centrifugation at 14,000 g for 15 min. Filters were then washed by the addition of 400 µL of UA buffer to the filter tops and centrifugation at 14,000 g for 15 min. This was followed by 2 further washes with 200 µL of UA buffer, and three washes with 50 mM Ammonium Bicarbonate (ABC buffer). The filters were then transferred to fresh collection tubes, and 50 µL of ABC buffer containing MS-grade Trypsin (Sigma) at 1:100 enzyme to total protein ratio was added directly to the filter tops, before being incubated at 37°C overnight in a humidified chamber to digest the proteins. Tryptic peptides were subsequently collected by centrifugation at 14,000 g for 10 min. This was combined with two further collections following the addition of 50 µL of ABC and 50 µL of 500 mM NaCl and centrifugation at 14,000 g for 10 min, respectively. The combined collected peptides were then acidified by the addition of STOP4 buffer (4% Acetonitrile, 1% Trifluoroacetic acid) at 1:1 (v:v) ratio.

An equivalent of 1 µg of digested peptide per sample was then loaded onto Evotip C18 tips (Evosep), according to manufacturer’s instructions, and subjected to mass spectrometry analysis with an Orbitrap Astral ZOOM (Thermo) coupled to an Evosep Eno LC system. Peptides were resolved on an EV1182 Evosep Performance Column, using a 200 Sample Per Day (200SPD) programme. MS was performed in Data Independent Acquisition (DIA) mode, according to(Guzman et al. 2026). Briefly, a full-MS resolution of 240,000, a scan range of 380 to 980 m/z, and a full-MS AGC target value of 5e6 was applied. Fragment ion scans were performed at 80,000 resolution, with a maximum Injection Time of 1.5 ms. A total of 300 windows of equal m/z widths were used for ion isolations. The isolated ions were fragmented using higher-energy collisional dissociation (HCD), with 25% normalized collision energy.

MS raw files were subsequently searched against an in silico generated spectral library from a *Drosophila melanogaster* proteome Fasta file (UniProt), using DIA-NN (version 2.2.0 Academia). Briefly, raw data files were searched using mostly default parameters; peptide length of 7-30 amino acids, precursor charge range of 1-4, precursor m/z range of 380 to 980 and fragment m/z range of 150 to 2000. Trypsin was selected as the protease, allowing for 2 missed cleavages. FDR was set at 1%, match between runs (MBR), unrelated runs and Protein Inference were enabled. The algorithm was set as 10.0 for mass accuracy, 4.0 for MS1 accuracy and default for all other parameters. The protein group output file from DIA-NN was loaded into Perseus (version 1.6.2.3) for Log2 transformation, filtering and statistical analysis.

### RT qPCR

RNA was Trizol (Sigma) extracted from five heads and thoraxes or abdomens from male adult flies and reverse transcribed as previously described (Koushika et al. 1999). Triplicate samples per cDNA were amplified by quantitative real-time PCR with a SensiFAST SYBR Hi-ROX Kit (BIO-92020, Bioline) under the following conditions: intial denaturation for 10 min at 95°C followed by 40 cyles (95 °C for 15 s, then 60 °C for 60 s). Analysis of CT values was done with the BioRad CFX Duet software. Differential expression was determined with the 2^−ΔΔCt^ method as described(Livak and Schmittgen 2001; Decio et al. 2021) using *ewg* amplified by primers F4 and R5 as reference gene(Haussmann et al. 2011).

### DNAzyme cleavage assay and Northern blotting

Total RNA was isolated from adult flies using Trizol (Sigma) and treated with DNase I (Ambion), phenol/CHCl_3_ extracted and ethanol precipitated in the presence of 1 µl glycogen (Roche). For the cleavage assay, 5 μg total RNA and 400 pmol of DNAzyme (each in a volume of 4 μl) were heated in separate tubes at 95 °C for 2 min followed by incubation at 25 °C for 10 min. To the DNAzyme, 8 μl of reaction buffer (200 mM KCl, 800 mM NaCl, 100 mM HEPES pH 7.0, 15 mM MgCl_2_, and 15 mM MnCl_2_) was added before adding the RNA and incubated at 25 °C for 2 h. Then RNA was extracted with phenol/CHCl_3_ extracted and ethanol precipitated in the presence of glycogen (Roche). The RNA was then separated on 1% agarose-formaldehyde gel, followed by performing a Northern blotting and methylene blue staining as described(Soller et al. 1999).

### Protein purification and and Electrophoretic Mobility Shift Assays

The *Drosophila* DM15 domain, full-length Sgt and the Sgt TPR domain (amino acids 116-217) were cloned into a modified pGEX vector containing a PreScission protease cleavage site and expressed in *E. coli BL21*(DE3) in LB media supplemented with 0.25 mM IPTG at 18°C overnight. After freezing, cells were resuspended in lysis buffer (PBS, 1 mM EDTA, 1 mM DTT, 1 mM PMSF). After sonication with a thin tip (20 s/40 s on/off for 3 min, Sonics), the resulting lysate was cleared by centrifugation in a Beckman Ultracentrifuge using an SW 41 T1 rotor swing-out rotor at 20,000 g and loaded onto Gluthathione Sepharose 4B custom-made column (Polyprep, BioRad) or GSTrap FF (Cytiva) and extensivly washed. Then the buffer was changed to cleavage buffer (50 mM Tris/HCl pH 7.5, 150 mM NaCl, 1 mM EDTA, 1 mM DTT, 0.025% NP-40). After washing, the protein was cleaved with procession protease overnight at 4°C and concentrated to 6 mg/ml using a centricon concentrator (Amicon) according to the manufacturer’s instructions.

EMSAs were done as previously described(McQuarrie and Soller 2024) using [α-32P]-GTP-capped oligonucleotides and 6% native polyacrylamide gels.

### Protein expression and purification for ITC experiments

The proteins for the recombinant expression of LARP1 DM15 and the tetratricopeptide repeat (TPR) of SGT described above were expressed in RosettaTM2 E. coli cells (NEB). Cells were cultured in Luria Broth medium at 37 °C, and expression was induced overnight at 18°C by addition of 0.5 mM isopropyl β-D-1-thiogalactopyranoside (IPTG). Cell pellets were resuspended in PBS pH 7,5% glycerol, 1 mM ethylenediaminetetraacetic acid (EDTA), 1 mM Dithiothreitol (DTT), one cOmplete™ tablet (Merck) per 50 mL of buffer, 2 mM phenylmethylsulfonyl fluoride (PMSF), 0.01 mg/mL DNAse I (Sigma) and 200 µg/mL lysozyme (Sigma), and lysed by sonication. The GST-tagged proteins were purified using high-performance GSTrap™ HP columns (Cytiva), eluting with a linear gradient from 0 to 20 mM of glutathione. The GST-tag was cleaved overnight with 5 µM in-house prepared 6xHis-PreScission protease(Ullah et al. 2016) in 50 mM TRIS-HCl pH 7.5, 150 mM NaCl, 1 mM EDTA, and 1 mM DTT at 4°C. The protease was removed by immobilised metal affinity chromatography (IMAC) using a HisTrap™ HP Column (Cytiva) equilibrated in 50 mM TRIS-Base pH 8, 300 mM NaCl, 10 mM Imidazole, 5% glycerol and eluted with a linear 0–100% gradient of 500 mM Imidazole. The cleaved GST-tag was then removed by several consecutive reverse GSTrap purifications. SGT TPR was additionally size excluded on a 16/600 Superdex 75 pg in storage buffer (20 mM TRIS-HCl pH 7.25, 100 mM KCl, 1 mM DTT). LARP1 DM15 was instead additionally cationic exchanged on HiTrapTM Heparin Sepharose FF (Cytiva) in 50 mM TRIS-HCl pH 6.25, 100 mM KCl, 1 mM DTT and applying a linear 0–50% gradient of 2 M KCl and dialysed in storage buffer. All protein samples were concentrated to ∼5 mg/mL using vivaspin centrifugal concentrators (Cytiva). Protein concentration was determined from the absorbance at 280 nm using the theoretical extinction coefficient calculated by ProtParam ExPASy(Duvaud et al. 2021). Protein purity was assessed in SDS-PAGE and nucleic acids contamination was assessed by measuring the 260/280 nm absorbance ratio. Samples in storage buffer were snap-frozen in liquid nitrogen and stored at -80°C.

### RNA preparation and handling

The RPL30 cap1/2 (m7Gppp(mC)(mU)UUUGCC), RPL30 cap1 (m7Gppp(mC)UUUUGCC), and RPL30 cap0 (m7GpppCUUUUGCC) RNA oligos were chemically synthesised by ChemGenes. The lyophilised powders were resuspended in DPEC treated water (SLS). The concentration was assessed by measuring the absorbance at 260 nm and using the extinction coefficient provided by the manufacturer.

### Isothermal titration calorimetry (ITC)

ITC experiments were performed using a MicroCal PEAQ-ITC instrument (Malvern Panalytical, Malvern, UK). Each experiment was conducted at 25°C and consisted of nineteen 2-μL injections of 2 seconds each, with a spacing of 120–150 seconds. The machine was set up to high-feedback mode, reference power 5 μcal/sec, and stirring speed of 700 rpm, and a 60-seconds pre-injection delay was applied for baseline stabilisation after equilibration. A summary of experimental conditions is reported in Table 1. Heat peaks were integrated and data fitted using the MicroCal PEAQ-ITC Analysis Software (Malvern). The fitting parameters were ΔH (reaction enthalpy change in kcal/mol), KD (equilibrium dissociation constant in M) and n (number of binding sites). The reaction entropy was automatically calculated using the relationships ΔG = −RT·ln(1/KD) (R 1.985 cal/mol·K, T 298 K) and ΔG = ΔH-TΔS. Experiments were repeated at least in three times. Buffer-into-buffer, protein-into-buffer and buffer-into-RNA control experiments were also performed.

**Table 1.** Summary of experimental conditions of the ITC experiments performed. The analyte was transferred in the adiabatic cell of the PEAQ-ITC and titrated with the titrant placed in the syringe. The syringe overall volume is 60 μL whereas the cell volume is 280 μL.

| **Titration – syringe into cell** | **[DM15] μM**<br>**in 60 μL** | **[RPL30] μM**<br>**in 280 μL** | **[STG**<br>**TPR] in**<br>**280 μL** |
| --- | --- | --- | --- |
| LARP1 DM15 into RPL30 cap0 | 150 | 15 | - |
| LARP1 DM15 into RPL30 cap1 | 150 | 15 | - |
| LARP1 DM15 into RPL30 cap12 | 50 | 5 | - |
| LARP1 DM15 into (RPL30 cap0 / SGT<br>TPR) | 150 | 15 | 20 |
| LARP1 DM15 into (RPL30 cap1 / SGT<br>TPR) | 150 | 15 | 20 |
| LARP1 DM15 into (RPL30 cap12 / SGT<br>TPR 20) | 50 | 5 | 20 |

### Cellular extracts and RNA stability assays

HeLa cell extracts were purchased (Ipracell, Belgium), and S2 cell extracts were made as described(Haussmann et al. 2022). RNA oligonucleotides were synthesised to harbour a 5’ triphosphate (Chemgenes) and capped with [α-32P]-GTP as described(Haussmann et al. 2022). 2’-*O*-ribose methylation was either included in the synthesis or added at the cap1 position by vaccinia CMTr VP39 (NEB). Oligos were then incubated in cellular extracts (40% v/v) under pre-mRNA processing conditions (1 mM ATP, 5 mM creatine phosphate, 2 mM MgAcetate, 20 mM KGlutamate, 1 mM, DTT, 20 U RNasin (Roche), and 5 μg/mL tRNA) on ice(Soller and White 2003). Reactions were quenched in 200 µl stop buffer (300 mM NaAc, pH 5.2, 10 mM EDTA, 1% SDS), phenol/CHCl_3_ extracted and ethanol precipitated in the presence of glycogen (Roche), separated on 12 % denaturing polyacrylamide gels, dried and exposed to a phosphoimager screen.

### UV-crosslinking assays, Western-blots, cap-binding protein purification and massspectrometry

For UV-crosslink assays [α-32P]-GTP capped oligonucleotides were incubated in cellular extracts (40% v/v, 10 µl total volume) under pre-mRNA processing conditions in microtiter plates (COSTAR 3894 V-bottom 96 well cell culture plates) as previously described(Soller and White 2003). After 15 min on ice, samples were illuminated with UV (265 nm) in a Strata linker for 15 min. The RNA was then digested with RNAse A/T1 mix (1 µl, Ambion) for 15 min. Then, 2X samples buffer (125 mM TrisHCL, pH 6.8, 4% SDS, 100 mM DTT, 15 % glycerol, 0.01 % bromphenol blue) was added, samples boiled, and proteins separated on 11.5% SDS-protein gels.

Immunoprecipitations and Western blots were done as previously described(Haussmann et al. 2016). Essentially, IPs were done in a final volume of 120 µL using 10 µL nuclear extract or UV cross-linking mix in NET 150 (150 mM NaCl, 0.5 M Tris-HCl at pH 7.5, 0.01% NP-40) at 4°C for 2 h with magnetic protein A/G beads (Pierce). After centrifugation, beads were washed with Pierce IP buffer. Proteins were separated on SDS-polyacrylamide gels, semi-dry blotted onto nitrocellulose and processed for detection with infrared-labelled secondary antibodies in a Licor. Antibodies used were anti-LARP1-Central 9347(Blagden et al. 2009), anti-LARP1 (AB86359, Abcam), anti-Sgt(Uytterhoeven et al. 2015)and anti-SGTA (11019-2-AP, Proteintech).

For purification of cap-binding proteins, biotin-pCp (Jena Bioscience) was ligated to a cold-capped RNA oligonucleotide with T4 RNA ligase 1 according to the manufacturer’s instructions (NEB) and upscaled for UV cross-linking (500 µl). Proteins bound to the oligo were then purified with streptavidin magnetic beads (Pierce) in Pierce IP buffer (25 mM Tris/HCl, pH7.4, 150 mM NaCl, 1 mM EDTA, 1% NP40, 5% glycerol), extensively washed and released by RNAse A/T1 digestion. Proteins were then separated on SDS-acrylamide gels, stained with Coomassie blue and subjected to protein determination of the cut bands by the University of Birmingham mass spectrometry facility.

### RNA in situ hybridizations, immunostainings and imaging

To analyze synapses at NMJs third instar wandering larvae were dissected in PBS and fixed with Bouin’s solution (Sigma-Aldrich, HT10132) for 5 min. The samples were washed three times in PBT (PBS with 0.1% TritonTM X-100 (Sigma, T8787) and 0.2% BSA) for 30 minutes. Antibody stainings were done as described(Haussmann et al. 2022) using the following primary antibodies: rat anti-HA (MAb3F10 1:20, Roche), rabbit anti-FLAG (M2, 1:250, SIGMA), rabbit anti-HRP (1:250, AffiniPure, Jackson Immuno Research), rabbit anti-Rpl35(Nyathi and Pool 2015) or DAPI (4’,6-diamidino-2-phenylindole,1 µg/ml) and stainings were carried out overnight at 4 °C followed by secondary antibodies (conjugated with Alexa Fluor 488 or Alexa Fluor 647 (1:500 Thermo Fisher Scientific) at RT for 2-3 h. NMJs were mounted in Vectashield (Vector Labs), then scanned with Leica TCS SP8, and images were processed and quantified using FIJI. For quantification of synapse staining, the comparative mean intensity of each genotype is normalized to wildtype.

Probes for RNA in situs were PCR amplified from cDNA and cloned into a modified pBluescript II SK+. Plasmids were linearised with Acc65I and used as templates for DIG-labelled antisense RNA synthesis with T3 RNA polymerase (Ambion). Transcription reactions (10 µl) contained 1 µg linearised DNA and DIG nucleotide labelling mix (Roche). Template DNA was removed with TURBO DNase (Ambion). The free nucleotides were removed with a G-50 MicroSpin columns (GE Healthcare) resulting in a final volume of labeled RNA in 50 µl DEPC-treaded water. Generally, 1 µl was added to 500 µl hybridization buffer, and further diluted if necessary.

RNA in situ hybridizations were done essentially done as described(Ustaoglu et al. 2021). Briefly, NMJ samples in PBT were incubated for 5 min in increasing amounts of 25%, 50% and 100% minimal hybridization buffer (50% formamide, 5× SSPE, 50 µg/ml heparin, 0.1% Tween-20), followed by 24 h at 42 °C in hybridisation buffer containing 50% formamide, 5× SSPE, 50 µg/ml heparin, 0.1% Tween-20, and 0.5 mg/ml denatured salmon sperm DNA and DIG-labelled probes (1:500). After removal of the hybridization buffer and washing with minimal hybridization buffer, samples were incubated in minimal hybridization buffer over night at 42 °C, followed by three 15-min washes in PBST. Hybridized probes were detected with sheep anti-DIG antibody (1:500; Roche) overnight at 4 °C and Alexa Fluor 546-conjugated secondary antibody overnight at 4 °C.

### Statistical analysis

All statistics in this study were done by GraphPad Prism 10. Two-tailed t tests were used for comparing two groups, and two-way ANOVA followed by a Fisher test was used for comparing multiple groups.

## Supplementary information

Supplementary materials include Figure S1-9.

### Authors’ contributions

MS conceived the project. MS directed the project. YWT performed genetic, biochemical and molecular biology experiments. YH generated CAGE-seq data, DWJM and FM carried out sequence analysis. NF and FPT performed neural activity tests. GA did biophysical experiments, SC analysed protein structures, MD, ELA and FM analysed proteins by mass spectrometry, YWT and MS wrote the manuscript. All authors read and approved the final manuscript.

## Acknowledgements

We thank P. Verstreken, Bloomington (NIH funding: P40 OD018537) and Kyoto stock centres for fly lines, the Manchester Fly Facility and S. Patel for fly injections, P. Verstreken for anti-Sgt, M. Pool for anti-RpL35 and D. Glover for anti-LARP1 antibodies, A. Berman for plasmids and communication of results before publication, W. Bian for help with cloning and Elise Cau for access to her incubator for the light-induction experiments..

## Funding

This work was supported by the Medical Research Council (MR/N013913/1) to D.W.J.M., the Biotechnology and Biological Science Research Council to M.S and Wellcome Trust to FM. G.A and M.R.C. acknowledge the Leverhulme Trust and the Biotechnology and Biological Science Research Council for funding.

### Availability of data and materials

All data generated or analysed during this study are included in the supplementary information files.

### Ethics approval and consent to participate

Not applicable.

### Consent for publication

Not applicable.

