## Supplementary material for "LARP1-DM15/Sgt is a reader for cap-adjacent 2’-*O*-ribose methylation in TOP ribosomal protein mRNAs required for localization to synapses": Suppl Figs

**for**

#### **Supplemental text**

**snoRNAs from down-regulated TOP mRNAs in *CMTr1/2<sup>null</sup>* mutants do not affect 2'-O-ribose methylation in rRNA**

Since levels of ribosomal proteins are tightly linked to rRNA levels (Lam et al. 2007), we compared 18S and 28S rRNA levels, which run as a single band on agarose gels (because 28S RNA in *Drosophila* is cleaved), between wild type and *CMTr1/2<sup>null</sup>* mutants, but did not find significant differences in rRNA amounts from the same number of individual flies corrected for their weight (Fig S2A). In addition, some downregulated TOP mRNA genes (*porthos*, *CG9253* and *RpL15*) contain snoRNAs. We compared 2'-O-ribose methylation at an unmethylated position G3253 in 28S rRNA, at position A469 in 18S methylated by the snoRNA encoded in *porthos* (CG9253) and at position C419 in 18S rRNA methylated by the snoRNA encoded in *RpL15* (Sklias et al. 2024) using a DNAzyme cleavage assay (Grzechnik et al. 2018), but did not find significant differences in *CMTR1/2<sup>null</sup>* mutants compared to wild type (Fig S2B and C).

### **DXO does not affect global levels of TOP ribosomal protein mRNAs**

In *C. elegans*, most transcripts (>80%) start with Gm added by a trans-spliced leader (Clemens et al. 2026). The remote DXO orthologue EOL-1 can suppress growth retardation and sterility of *CMTr1* mutants (Clemens et al. 2026). To test whether DXO destabilizes TOP mRNAs in *Drosophila* we obtained a transposon insertion mutant in the beginning of the *DXO* open reading frame likely constituting a complete null mutant (*DXO<sup>null</sup>*, *DXO* in flies is called *Rail*). Like *CMTr1/2<sup>null</sup>* mutants, *DXO<sup>null</sup>* mutants are viable as well, and DXO is expressed broadly with an enrichment in the brain (Fig S4A and B).

Removal of *DXO* in *CMTr12<sup>null</sup>* mutants significantly increased expression levels of TOP ribosomal mRNAs, but removal of DXO alone did not increase them further (Fig S4C).

At an organismal level, body weight of *CMTr1/2<sup>null</sup>* mutants is about 20% smaller than the weight of control flies. Removal of *DXO* in *CMTr1/2<sup>null</sup>* flies restores body weight to wild type levels and *DXO* mutants have a significantly increased body weight compared to control flies (Fig S4D and E).

### **LARP1 proteins stabilize TOP ribosomal protein mRNAs in *Drosophila***

La related protein family (LARP) proteins contain a highly conserved La-module consisting of a La-motif and an RNA recognition motif (RRM) (Figure S4A), which has key roles in TOP mRNA regulation (Berman et al. 2021). In *Drosophila* there are three main *LARP* genes (*LARP1*, *LARP4B* and *LARP6*) with orthologues in humans (*LARP1*, *LARP4B* and *LARP6*) (Fig S5A). The most highly expressed LARP family protein during development in *Drosophila* is *LARP1* and expression is ubiquitous with an enrichment in the brain (Fig S5B and C) (Blagden et al. 2009; Burrows et al. 2010). Expression of *LARP4B* and *LARP6* is lower and more tissue-restricted. To examine whether these threeLARPs impact on TOP mRNA expression levels, we obtained null mutants, which were viable and fertile as trans-heterozygotes of two alleles (*LARP1*<sup>43</sup>/*LARP1*<sup>MI15379</sup>, *LARP4B*<sup>CR70229-TG4.0</sup>/*Df(3L)ED4293* and *LARP6*<sup>KG04043</sup>/*Df(2R)ED2354*), except for *LARP1*<sup>43</sup>/*LARP1*<sup>MI15379</sup>, which were sterile.

The combination of *LARP1* and *LARP6* mutant was lethal (64 %, n=390), whereas mutant combinations of *LARP1* with *LARP4B* (41%, n=355) and *LARP4B* with *LARP6* (11%, n=498) remained viable, suggesting partially redundant functions between *LARP1* and *LARP6*.

We then determined expression levels of *RpL3,5,35* and *RpS16* genes by qPCR in these *Larp* mutants. In all of them expression levels were strongly reduced, more than in *CMTTr1/2*<sup>null</sup> mutants indicating additional functions of LARP1 in TOP mRNA turn-over beyond protection through the cap. In *LARP6* and *LARP4B* mutants, expression of these ribosomal protein genes was less reduced compared to *LARP1* (Fig S5D).

#### **Loss of *DXO* increases LARP1 DM15, but not Sgt at synapses**

Although loss of the decapping enzyme *DXO* did not result in increased levels of TOP ribosomal protein mRNA in *CMTTr1/2*<sup>null</sup> flies (Fig S4A), specifically the pool of TOP ribosomal protein mRNAs localizing to synapse could be affected. In the absence of *DXO*, levels of DM15, but not Sgt were higher at synapses (Fig S9B and C). In the absence of cOMe and *DXO*, DM15 also did not localize to synapse, while Sgt was unaffected (Fig S9B and C). We noted however that expression of DM15 resulted in lethality of *DXO*<sup>null</sup> *CMTTr1/2*<sup>null</sup> flies.

### Supplemental figures

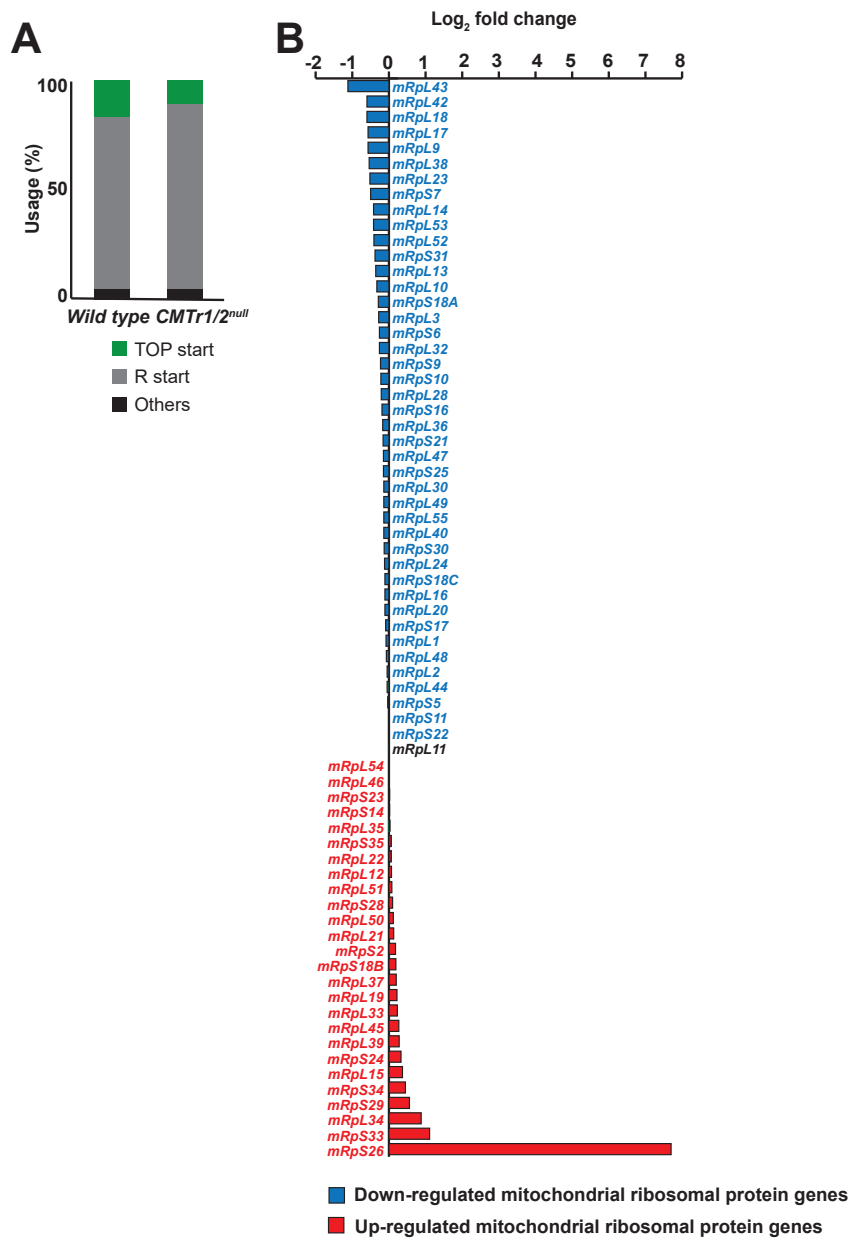

**Figure S1. Transcription start site dependence of expression.**

A) Comparison of transcription start site use (first transcribed nucleotide) in wild type and *CMTr1/2<sup>null</sup>* mutant flies.

B) Expression differences (log<sub>2</sub> fold changes) mitochondrial ribosomal proteins between wild type and *CMTr1/2<sup>null</sup>* mutant flies from CAGE-seq (blue and red bars: down- and up-regulated).

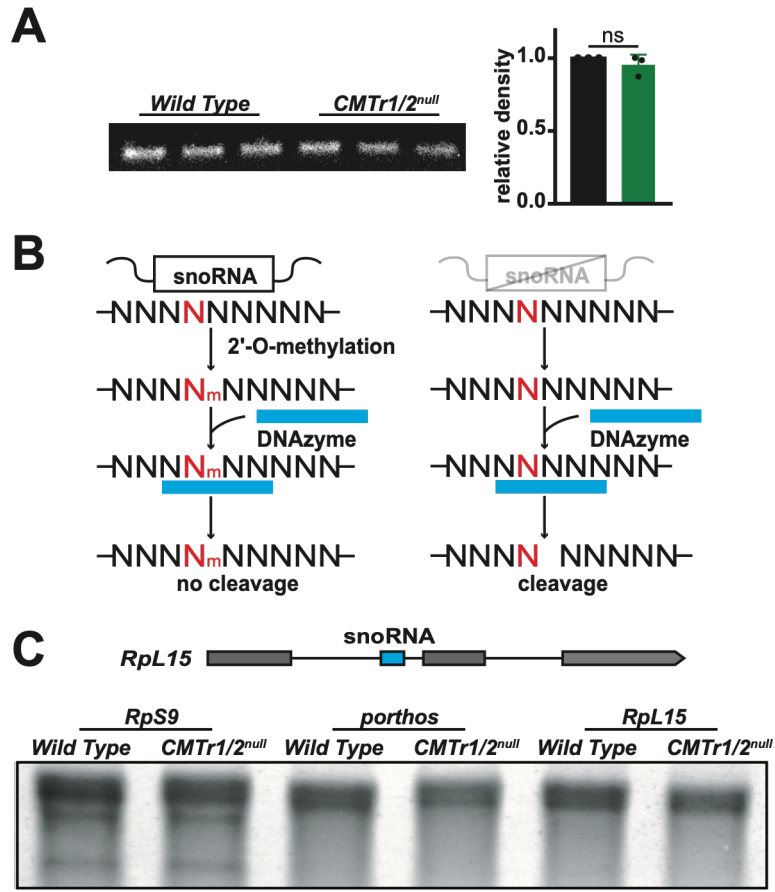

**Figure S2. The absence of cOMe does not affect rRNA amount and methylation levels.**

A) Denaturing agarose gel depicting rRNA from five wild type and *CMTr1/2<sup>null</sup>* male flies and quantification showing the mean (n=3) with the standard error corrected for the weight of flies.

B) Schematic depiction of 2'-O-ribose methylation detection by a DNAzyme cleavage assay.

C) Schematic of the *RpL15* gene (top) depicting the exon-intron structure and the position of the snoRNA and methylene blue stained Northern blot of DNAenzyme treated total RNA of wild type and *CMTr1/2<sup>null</sup>* flies for an unmethylated position in 28S rRNA (left), for position A469 in 18S methylated by the snoRNA encoded in down regulated *porthos* gene (middle) and for position C419 in 18S rRNA methylated by the snoRNA encoded in down regulated *RpL15* gene (right). Asterisks: cleaved fragments.

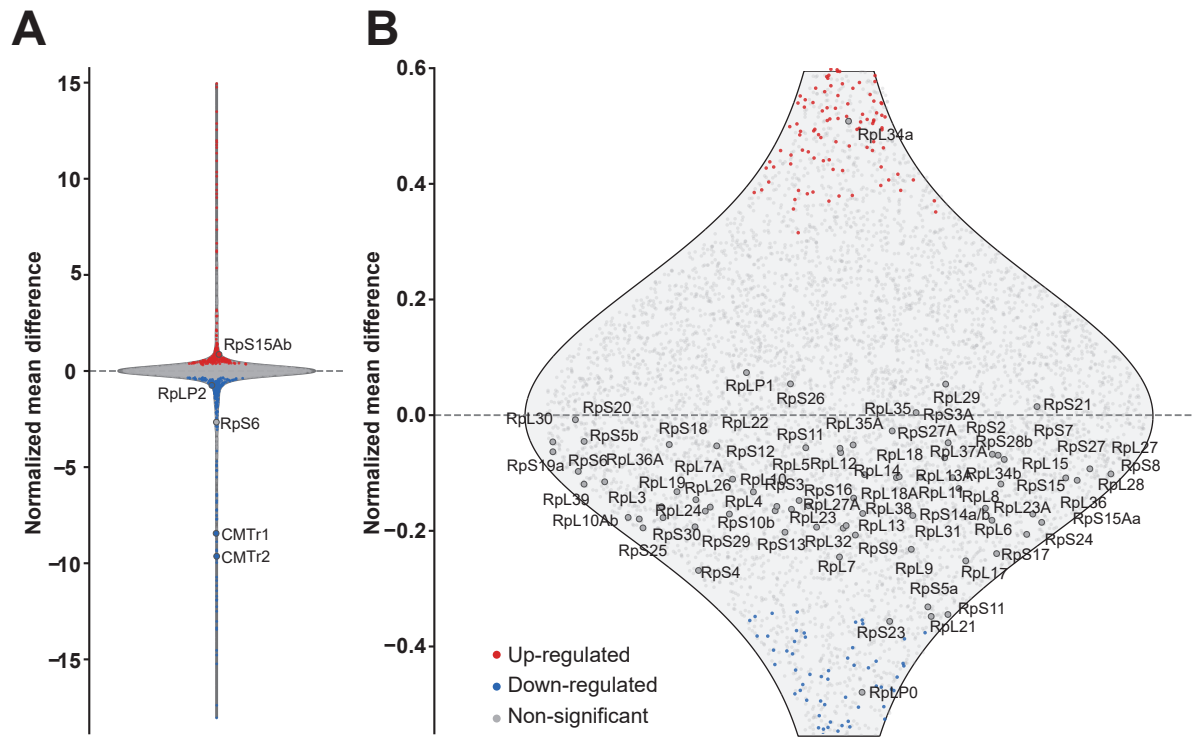

**Figure S3. Protein analysis by mass spectrometry.**

A-B) Violin plot of all proteins (A) and TOP ribosomal proteins (B) differentially expressed in head/thorax of wild type compared to *CMTr1/2<sup>null</sup>* mutants. Upregulated proteins are shown as red dots, and downregulated proteins as blue dots. Non-significant proteins are indicated with grey dots.

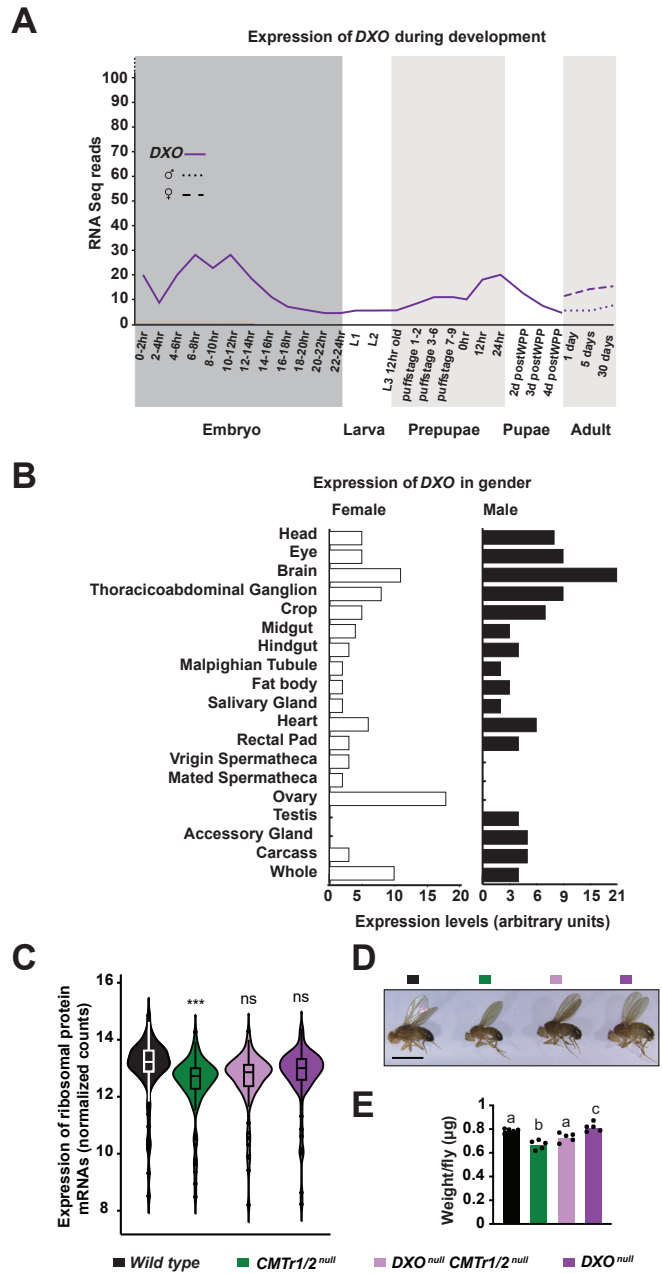

**Figure S4. *Drosophila* *DXO* is broadly expressed.**

A and B) *DXO* expression profile during development from RNAseq (A) or in various tissues from microarrays (B) (from flybase.org).

C-E) Expression levels of TOP ribosomal protein mRNAs in neuron-enriched head/thorax (C), pictures illustrating the size of flies (D) and weight (E) of wild type (black) and *CMTr1/2*<sup>null</sup> (green), *DXO*<sup>null</sup> *CMTr1/2*<sup>null</sup> (light purple) and *DXO*<sup>null</sup> (dark purple) mutants. The weight is shown as mean (n=5, 50 flies per replicate) with the standard error. Statistically significant differences are indicated by different letters ($P \leq 0.0001$ except $p \leq 0.005$ for c).

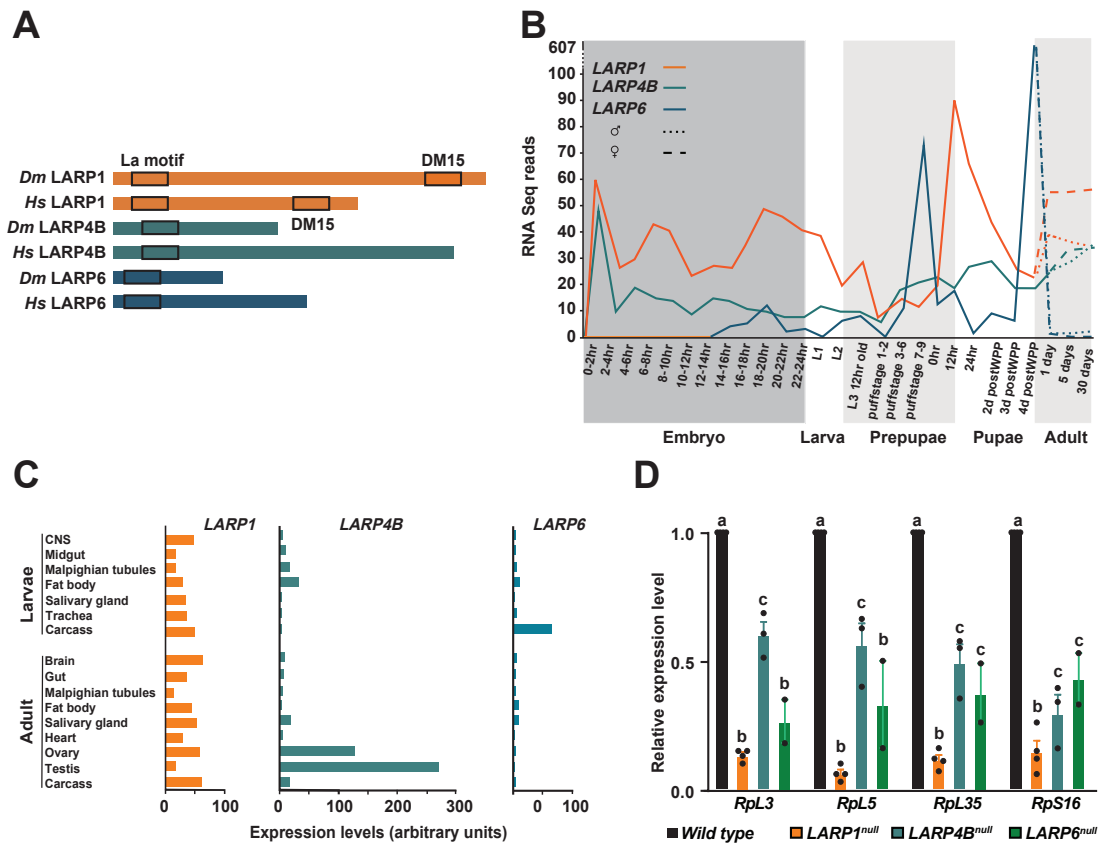

**Figure S5. Ribosomal protein genes are downregulated in *LARP1*, *LARP4* and *LARP6* mutants.**

A) Schematic protein structure of *Drosophila* LARP1 (orange), LARP4B (teal) and LARP6 (blue), and human LARP1, LARP4B and LARP6 depicting La and DM15 domains.

B and C) *LARP1*, *LARP4B* and *LARP6* expression profile during development from RNA-seq (B) or in various tissues from microarrays (C) (from flybase.org).

D) Expression levels of *RpL3*, *RpL5*, *RpL35* and *RpS16* genes determined by RT-qPCR are shown as mean (n=3-4) with the standard error for wild type (black), *LARP1*<sup>null</sup> (orange), *LARP4B*<sup>null</sup> (teal) and *LARP6*<sup>null</sup> (blue). Statistically significant differences from ANOVA post-hoc comparisons are indicated by different letters ($p \leq 0.001$ except $p \leq 0.027$ for c).

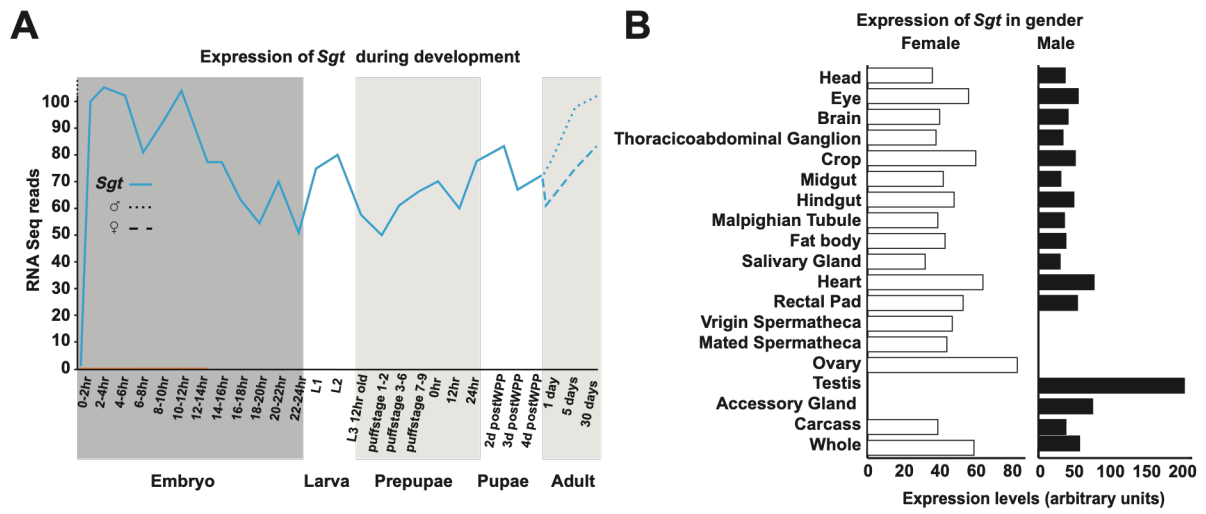

**Figure S6. *Drosophila* *Sgt* expression profile.**

A and B) *Sgt* expression profile during development from RNAseq (A) or in various tissues from microarrays (B) (from flybase.org).

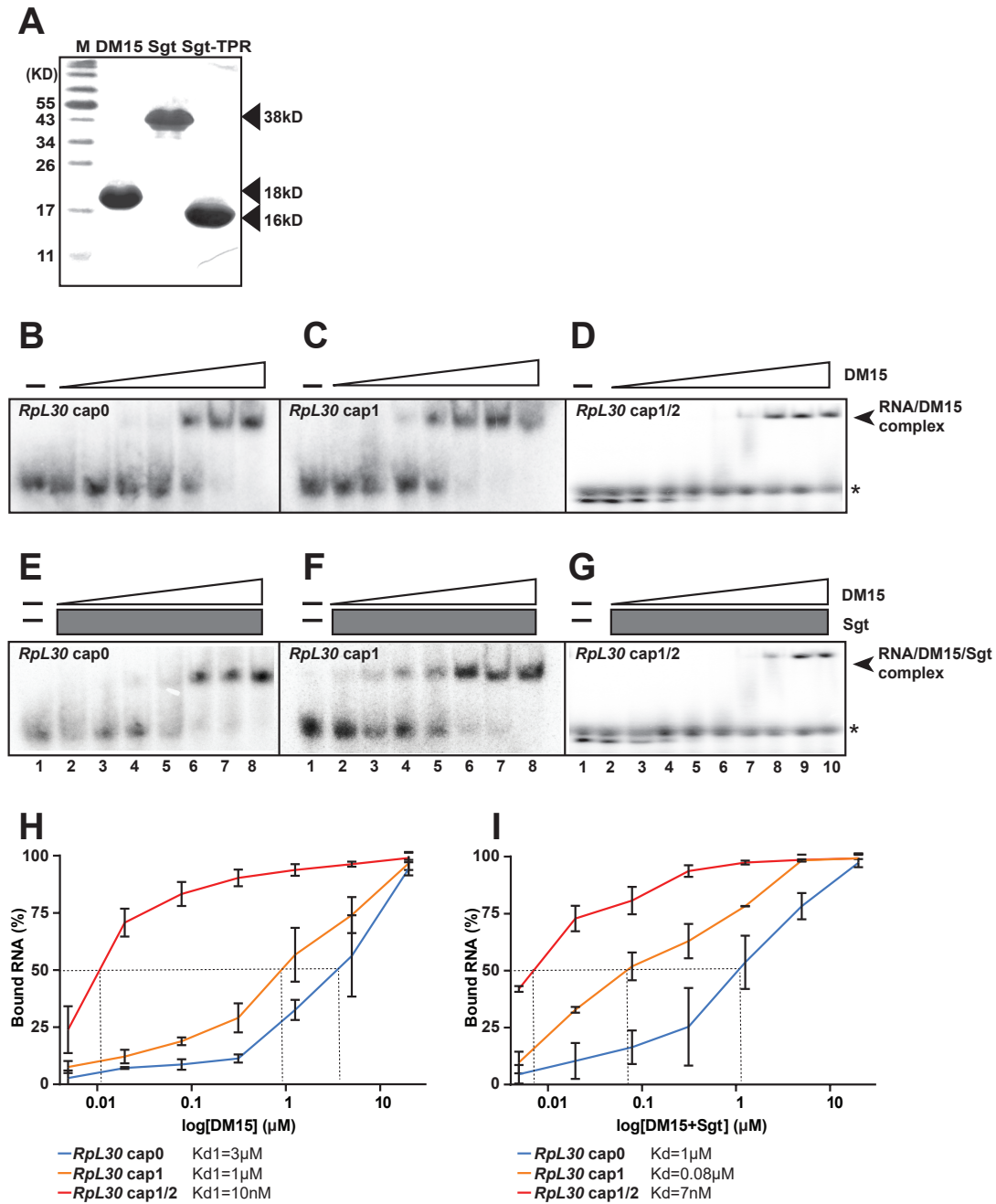

**Figure S7. Binding of LARP1-DM15 to TOP RNA is enhanced by cOMe and Sgt.**

A) SDS-gel of purified proteins for DM15, SGT and SGT-TPR domain.

B-G) EMSAs of various concentrations of DM15 (start from 20 μM, 4-fold serial dilution) and RpL30 cap0 (32 nt, 1 pM, B), cap1 (C) and cap1/2 (D) alone or in the presence of Sgt (20 μM, E-G) and quantification thereof (H and I, cap0 in blue, cap1 in yellow, cap1/2 in red). Means (N=3) with standard error are shown with Kd values at the bottom. The asterisk in G denotes a background contaminant from use of an eight nucleotide RNA oligo in this experiment.

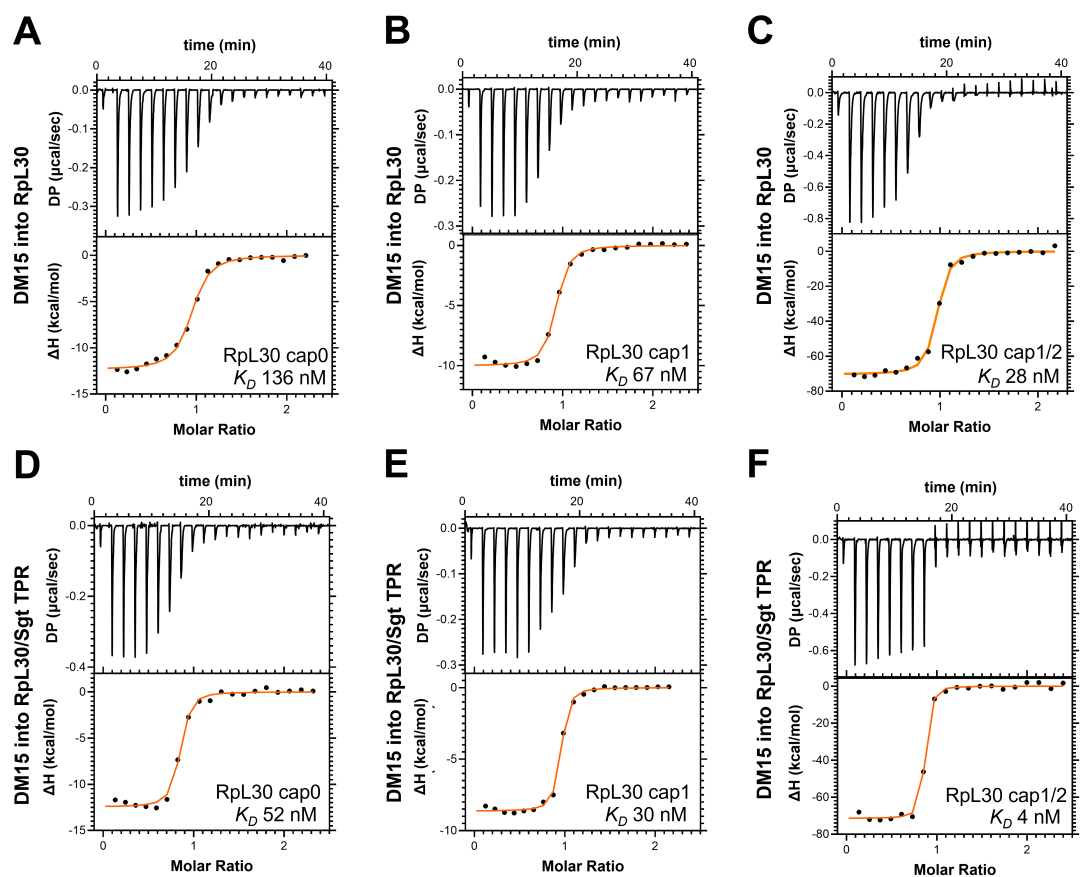

**Figure S8. ITC binding analysis of LARP1-DM15 to TOP RNA enhanced by cOMe and the Sgt TPR domain.**

A-F) Representative ITC thermograms (top panels) and data fitting (bottom panel) of the LARP1-DM15 titrations into Rpl30 (8 nt, 1 pM) cap0 (A and D), cap1 (B and E) and cap1/2 (C and F). D-F) in the absence (A-C) or presence of the Sgt TPR domain (D-F).

Table (below) with thermodynamic parameters obtained in ITC experiments.

| Titration: Syringe component/cell component | $K_D$ (nM) | N (binding sites) | $\Delta H$ (kcal/mol) | $\Delta G$ (kcal/mol) | $-T\Delta S$ (kcal/mol) |
| --- | --- | --- | --- | --- | --- |
| dLaRP1 DM15 / RPL30 cap0 | $136 \pm 4.4$ | $0.83 \pm 0.07$ | $-15.4 \pm 1.3$ | $-9.4 \pm 0.02$ | $6.0 \pm 1.3$ |
| dLaRP1 DM15 / RPL30 cap1 | $66.5 \pm 1.5$ | $0.81 \pm 0.02$ | $-9.52 \pm 0.2$ | $-9.8 \pm 0.01$ | $-2.6 \pm 0.2$ |
| dLaRP1 DM15 / RPL30 cap1/2 | $27.5 \pm 2$ | $0.82 \pm 0.04$ | $-67.5 \pm 2.6$ | $-43.2 \pm 0.2$ | $22.5 \pm 2.2$ |
| dLaRP1 DM15 / RPL30 cap0 and dSGT TPR | $52.4 \pm 3.2$ | $0.85 \pm 0.02$ | $-13.3 \pm 0.5$ | $-9.9 \pm 0.03$ | $-3.3 \pm 0.6$ |
| dLaRP1 DM15 / RPL30 cap1 and dSGT TPR | $30.0 \pm 1.9$ | $0.79 \pm 0.04$ | $-10.1 \pm 0.5$ | $-10.3 \pm 0.04$ | $-1.2 \pm 0.6$ |
| dLaRP1 DM15 / RPL30 cap1/2 and dSGT TPR | $3.8 \pm 0.1$ | $0.84 \pm 0.02$ | $-65.2 \pm 2.3$ | $-48.1 \pm 0.1$ | $17.1 \pm 2.2$ |

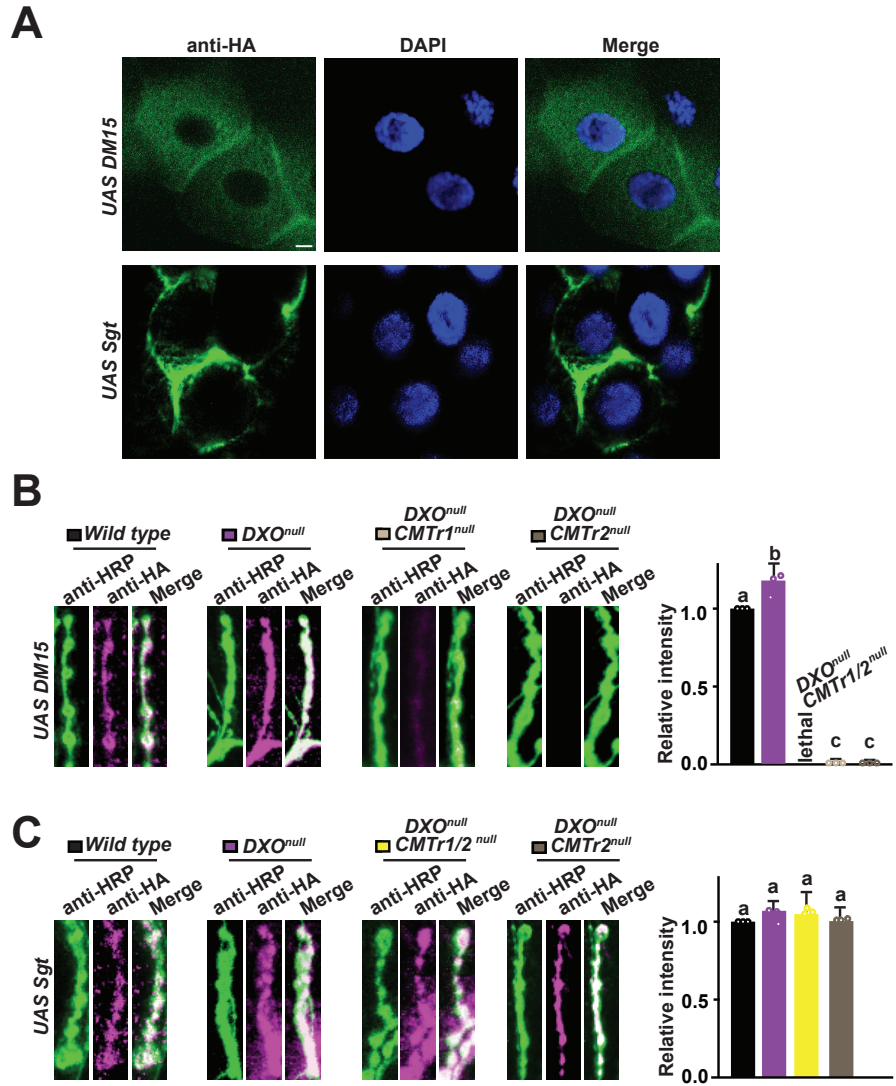

**Figure S9. Cellular localization of LARP1-DM15 and Sgt expressed in salivary glands and *DXO* localization dependence at synapses.**

A) Salivary gland cells of UAS HA DM15 (top) or UAS HA Sgt (bottom) expressed with *elavGAL4<sup>c155</sup>* stained with anti-HA antibodies (green) and DAPI (nuclei in blue).

B and C) Representative images from staining of synapses at third instar NMJs pre-synaptically expressing *UAS HA DM15* or *UAS Sgt* with *elavGAL4<sup>c155</sup>* in wild type (black), *DXO<sup>null</sup>* (purple), *DXO<sup>null</sup> CMT1/2<sup>null</sup>* (yellow), *DXO<sup>null</sup> CMT1<sup>null</sup>* (light brown) and *DXO<sup>null</sup> CMT2<sup>null</sup>* (dark brown) stained with an anti-HA antibody (magenta) and anti-HRP-antibodies (green) and quantification of staining intensity showing the mean with standard error in arbitrary units. Note that *elavGAL4<sup>c155</sup> UAS HA DM15* expression in *DXO<sup>null</sup> CMT1/2<sup>null</sup>* mutants is lethal.
